# Glow with the Flow: Reproducible Analysis of Transiently Transformed Protoplasts Using Dual Fluorescent Reporters in R

**DOI:** 10.1101/2025.07.26.666912

**Authors:** Joseph S. Taylor, Elyse A. Shoppell, Bastiaan O. R. Bargmann, R. Clay Wright

## Abstract

Transient transformation assays using protoplasts have become a widely employed technique in plant research. Positive fluorescent selection was subsequently developed to assess the effect of transient effector gene expression in only successfully transfected cells using flow cytometry. This process, though effective, often requires considerable manual effort and subjective judgment to quantify reporter gene expression in the intended cell populations. To address this, we introduce a new, open-source workflow based on the R programming language. This method enhances the reproducibility and scalability of such experiments, which enable rapid study of gene regulation and signal transduction in plants. This workflow is available at https://github.com/PlantSynBioLab/positive-fluorescence-selection.

## Introduction

Transient transformation of protoplasts is a widely used technique in plant research due to its relative ease and utility (Bargmann and Birnbaum 2009; Nanjareddy et al. 2016; Lin et al. 2018). An advantage of this technique is the speed and throughput with which experiments can be conducted in addition to enabling quantitative measurements of reporter-gene activation. These assays are often used to quantify signaling and gene regulatory networks by assessing gene expression, promoter elements, transcription factor or signal-transduction protein activity, and subcellular protein localization (Tiwari et al. 2006). Although transient assays in protoplasts to assess reporter-gene activation have proved invaluable in elucidating the elements involved in the regulation of gene expression, the approach comes with limitations. Historically, transcription factor studies using protoplasts involved co-transfection with up to three plasmids: 1) a promoter::reporter construct (e.g., *β-GLUCURONIDASE* or firefly *LUCIFERASE*); 2) an effector construct (e.g., a transcriptional activator/repressor); and 3) a constitutively expressed transformation reporter (e.g., Renilla *LUCIFERASE*/*RLuc*) (Liu et al. 1994, Ulmasov et al. 1997ab). Spectroscopy of bulk cultures was used to measure reporter expression. These initial systems had drawbacks, including skewed measurements from variable protoplast transformation efficiency and the reporter signal being detected in cells lacking the effector. These assays were later adapted to use a genome-integrated reporter system, forgoing the need for triple co-transfection, reducing experiment complexity. However, the field still lacked the ability to selectively measure only successfully transformed cells, which typically represent a subset of the total population (Tiwari et al. 2006).

Positive fluorescent selection of protoplasts was developed as a contemporary approach to quantify transient effector gene expression in only successfully transformed cells via flow cytometry (Bargmann and Birnbaum 2009). This system allows for analysis of only the transformed cell population, requiring just a single vector input coding for the effector (pBeaconRFP). In addition to the effector, the vector also encodes a fluorescent protein reporter used for identification of transformed cells (e.g., red fluorescent protein, RFP). The activation of a complementary genome-integrated fluorescent reporter (e.g., green fluorescent protein, GFP) is then measured in only those RFP-positive cells expressing the transfected effector gene. Furthermore, the addition of fluorescence-activated cell sorting of the successfully transformed cell population enables additional downstream analyses (e.g., transcriptome analysis via RNA sequencing).

Flow cytometry paired with positive fluorescent selection enables rapid, single-cell analysis of transient expression in protoplasts (Bargmann and Birnbaum 2009; Alvarez et al. 2020; Beard et al 2021; Gonzalez et al. 2021). However, reproducibility of data analysis is crucial (Sandve et al. 2013). In typical implementations, proprietary flow cytometry analysis software is used to identify the transformed-cell population and quantify reporter gene expression (FlowJo Software 2023; Sony Corporation 2015; ThermoFisher Scientific 2015). Additionally, the identification of viable, successfully transformed cells using standard flow cytometry software is time-consuming and relies heavily on subjective gating decisions (Bio-Rad 2023).

To address these issues, we developed an open-source, data-driven analysis workflow using the R programming language to improve the reproducibility and scalability of transient expression assays for investigating reporter gene activation. The positive fluorescence selection workflow can be broken down into a series of steps: 1) Annotate, compensate, and transform the dataset, 2) distinguish individual cells from bubbles, debris, and doublets, 3) identify successfully transformed (e.g., RFP-positive) cells, 4) measure reporter activation (e.g., *promoter::GFP*) in transformed cells, 5) conduct statistical analysis, and 6) generate figures. We present an example vignette using this workflow to standardize and semi-automate analyses for high-throughput (e.g., 96-well plate) transient expression experiments using dual-fluorescent reporter systems (e.g., GFP and RFP).

We have assembled this workflow using the core flow cytometry infrastructure of the Bioconductor R package repository (Hahne et al. 2009; 2025; Finak et al. 2014; Wright et al. 2025; Van et al. 2009; 2015). While R programming requires the use of a text/command-line-based interface, it also offers the ability to develop templated reports with integrated descriptive text, R code, and figures. This format, known as R-markdown, or more recently Quarto, provides an ideal medium for transparently communicating the complete analysis and results in a single, easily reproducible format (Allaire et al. 2023; Xie et al. 2023) Additionally, we have found the community of R users to be very welcoming and helpful to new learners, as evidenced by the numerous, free, community-developed resources for learning (find a short list of excellent resources at https://education.rstudio.com/learn/beginner/). The included vignette uses the R-markdown format where each step of data processing and analysis is described in greater detail. For much of this workflow, we utilize the flowCore (Hahne et al. 2009), flowStats (Hahne et al. 2025), and flowtime (Wright et al. 2025) packages from Bioconductor. To define data-driven gates we apply clustering algorithms as implemented in the Bioconductor packages and openCyto (Finak et al. 2014). This allows us to define rectangular or elliptical gates which contain a certain quantile of a specified population. These algorithmically defined gates are then plotted using the ggcyto package (Van et al. 2009; 2015) for visual inspection and potential adjustment of the quantile parameter.

Ultimately, this workflow enables a fully transparent, open source, and reproducible data analysis pipeline for transient expression assays, adaptable to various flow cytometry platforms. Annotation steps included in the workflow facilitate attachment of experimental metadata to the dataset according to FAIR principles (Wilkinson et al. 2016) as required by FlowRepository (Spidlen et al. 2012), a central repository of flow cytometry datasets. We have validated this workflow with the included example dataset and demonstrated its utility in multiple peer-reviewed publications (Gonzalez et al. 2021; Taylor et al. 2025).

While here we focus on experiments using plant protoplasts, this workflow is applicable to any cell type including yeast, mammalian cells, and fungal protoplasts.

## Methods

### Example dataset overview

The worked example data consists of *Arabidopsis thaliana* root protoplasts carrying an integrated green-fluorescent auxin reporter (*DR5rev::GFP*; ABRC stock CS9361 (Benková et al., 2003)). These cells were transfected with the pBeaconRFP vector (Bargmann and Birnbaum 2009), which encodes RFP and an effector gene–either *AUXIN RESPONSE FACTOR 7/NON-PHOTOTROPIC HYPOCOTYL 4 (ARF7/NPH4)* or *β-GLUCURONIDASE* (*GUS*, an inert control)–or were mock-transformed. Additionally, a subset of the cells transfected with the *GUS* construct were also treated with 100 nM indole-3-acetic acid (auxin) to induce expression of *DR5rev::GFP*.

### Transient expression assay and flow cytometry

Described similarly in Gonzalez et al. (2021) and Bargmann and Birnbaum (2009), roots of one-week-old seedlings were harvested and placed into a 500 mL flask, gently shaking at approximately 75 RPM and 28 ºC in 50 mL enzymolysis solution (1.25% [w/v] Cellulase R-10 [Yakult Pharmaceutical Industry Co., Japan], 0.3% [w/v] Macerozyme R-10 [Yakult Pharmaceutical Industry Co., Japan], 0.4 M mannitol, 20 mM MES, 20 mM KCl, 10 mM CaCl2, 0.1% [w/v] bovine serum albumin; pH was adjusted to 5.7 with KOH pH 7.5) for 5 h. The protoplast solution was filtered through a 40 µm Falcon cell strainer (VWR, Radnor, PA, USA), divided over 15 mL conical tubes, and centrifuged for 5 min at 500 g. Cells were gently washed in the enzymolysis buffer (no enzymes added), centrifuged for 5 min at 500 g, resuspended in 1 mL of the enzymolysis buffer, and then counted with a hemocytometer. Cells were then washed with transfection solution (0.4 M mannitol, 15 mM MgCl2 hexahydrate, 4 mM MES; pH was adjusted to 5.7 with KOH), centrifuged again, and resuspended in transfection solution with a final density of 4*10^6^ protoplasts/mL. Conical tubes (15 mL) were prepared for each transfection with 25 µg of plasmid DNA (1 µg/µL) and 250 µL of protoplasts in transfection solution. Protoplasts were transformed with the pBeaconRFP vector carrying *ARF7/NPH4, GUS*, or not transfected. Next, 250 µL PEG solution (40% [w/v] PEG 1500, 0.2 M mannitol, 0.1 M CaCl2) was added, and the suspension was mixed by flicking the tube repeatedly. Suspensions were immediately washed with 15 mL of enzymolysis buffer, centrifuged, and resuspended in 750 µL enzymolysis buffer. One of the two GUS samples was treated with 100 nM indole-3-acetic acid. Protoplast suspensions were then divided three ways in a 96-well plate (250 µL per well) to generate three technical (measurement) replicates. The plate was incubated overnight in the dark at room temperature.

The GFP intensity in RFP-positive cells was quantified using a Sony SH800S fluorescence activated cell sorter with 488 nm excitation and 525/50 nm emission for GFP and 561 nm excitation and 617/30 nm emission for RFP. Data was exported from the proprietary cytometry software in the flow cytometry standard file format (.FCS).

Data was processed as described below and R markdown script was developed and tested using R version 4.4.3. Visualization we performed using R packages ggcyto and ggplot2 (Van 2018; Wickham 2016).

## Workflow

The full analysis workflow we describe below is available at https://github.com/PlantSynBioLab/positive-fluorescence-selection, as well as in the Supplemental Materials.

### Importing and annotating the dataset

Importing the FCS data as a flowSet object in the R environment using the provided workflow allows for annotating, compensating, and converting the flow cytometry files into a format which can be used to calculate summary statistics and generate publication-ready figures.

Annotations are important for enabling repeatable and/or novel analyses by independent researchers. If the annotation meets or exceeds the minimal information standards for flow cytometry experiments (MIFlowCyt) defined by Lee et al. (2008), the dataset can be deposited into FlowRepository (Spidlen et al. 2012) to be shared with others in the scientific community. Briefly, the MIFlowCyt standard makes recommendations about descriptions of relevant specimens and reagents, the configuration of the instrument used to perform the assays, and the data processing steps taken to interpret the primary data. Annotations should be specific, including information such as sample name, expression vector, treatment type, duration, concentration, etc. Annotated flowSets can be exported, allowing others to reload the dataset with experiment information preserved for future use.

### Visualizing raw data

We initially check event distributions of each sample on a log scale to screen for any inconsistencies, possibly caused by contamination or mechanical issues. This is done by generating forward scatter area (FSC-A; event size) versus side scatter area (SSC-A; internal complexity) dot plots (Figure 1; Kazovsky 1984). Log transformation of the data is useful to address skewness and achieve a near-normal distribution (West 2021; Hammill 2021). FSC and SSC area is used instead of height because it provides more reliable measures of these parameters as the latter solely measures the peak signal. Issues at this step may manifest as inconsistent clusters of events between samples in which case the compromised sample could be removed. Debris typically will exhibit lower relative levels of FSC and SSC and is usually found in the bottom left of density plots (BIO-RAD 2025). Given that all samples in the provided flowSet are from the same source tissue and free of contamination, each sample distribution will appear relatively similar (Figure 1).

**Figure 1:**
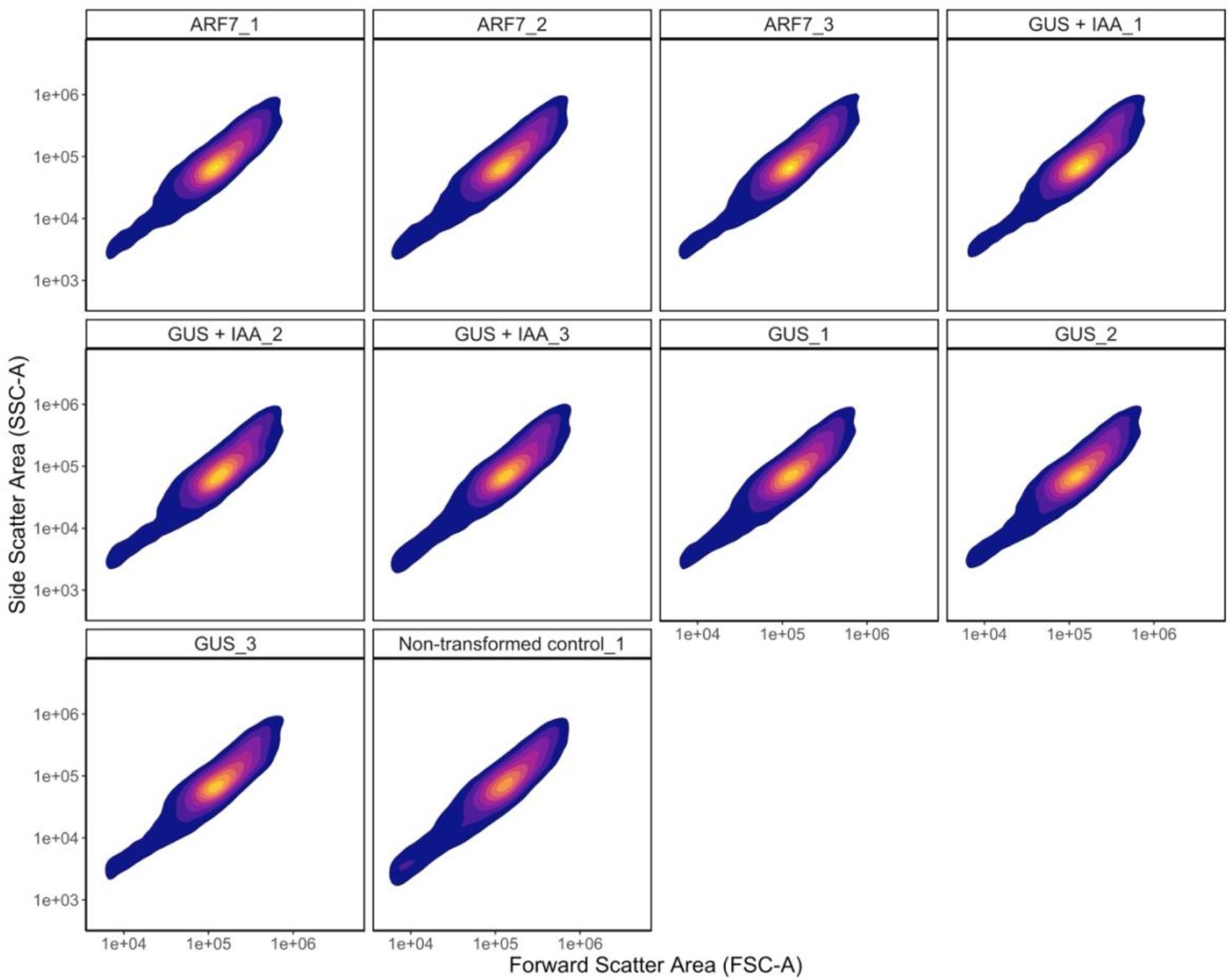
Log scale density gradient of all events on a forward scatter area (FSC-A) by side scatter area (SSC-A) distribution of all events for each sample in the flowSet. Lighter colors represent higher density, and darker colors represent lower density.

### Compensating the dataset

The use of multiple fluorescent reporters can lead to spillover, which may artificially inflate fluorescent signal measurements (Ortolani 2022). Spillover happens when fluorescent reporters have overlapping emission spectra. For example, in our flowSet, GFP is measured in the first fluorescent light channel (FL1; green fluorescence detector), but some signal also leaks into the FL3 channel (red fluorescence detector), where RFP is measured. Therefore, without correction, cells expressing high levels of GFP will have artificially inflated RFP measurements, biasing further analyses (Figure 2). Compensation corrects for fluorescence spillover by subtracting the proportion of signal that bleeds into non-primary detectors, allowing accurate measurement of each fluorophore’s true signal (Ortolani 2022). In some cases, the emission spectra and measurement channels of the two fluorescent proteins may not result in any spillover and thus compensation is not required. To determine the amount of spillover and calculate a compensation matrix, control cells that express each fluorescent protein individually as well as those with no fluorescent proteins are required (Ortolani 2022). Ideally, these controls should be included in every experiment to ensure compensation is calculated accurately. Spillover can be visualized in the flowSet by generating a GFP intensity vs RFP intensity dot plot of all events before and after applying the compensation matrix calculated by the SH800s flow cytometer (Figure 2; Sony Corporation 2015). The plots reveal three distinct event populations: those with high RFP and GFP, high GFP only, and low levels of both signals. In the uncompensated flowSet, as GFP signal increases, so does the RFP signal. Consequently, gating of high RFP events risk the inclusion untransformed cells with inflated GFP measurements in downstream analyses. In contrast, the compensated flowSet shows a much clearer separation between events high in both GFP and RFP and those high in GFP signal alone (Figure 2).

**Figure 2:**
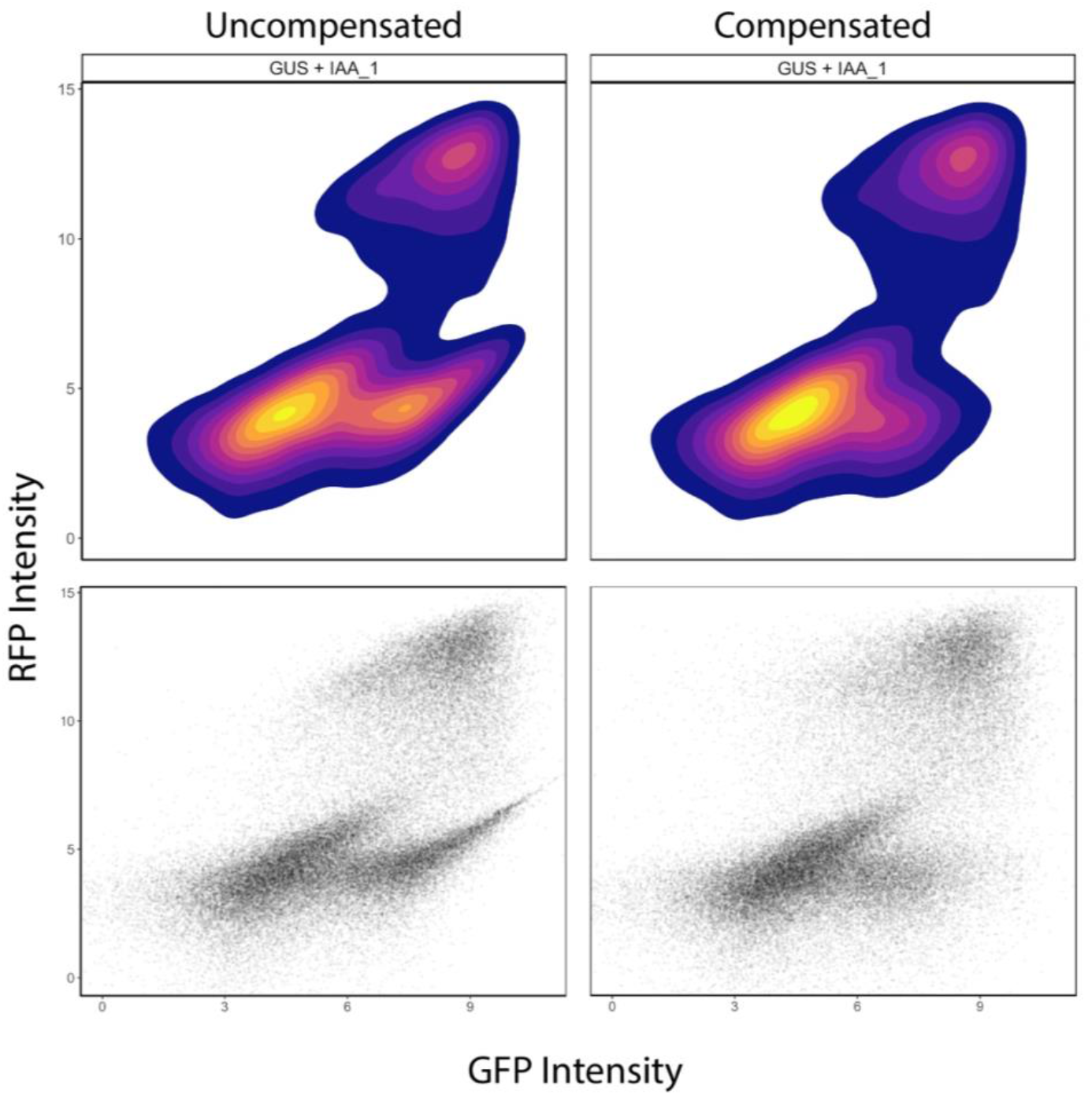
Log scale comparison of uncompensated **(left)** to compensated **(right)** GFP-area by RFP-area distribution of all events (arbitrary units). Shown in density **(top)** and dot **(bottom)** plot formats. Lighter colors represent higher density, and darker colors represent lower density.

### Defining a live-cell gate based on positive RFP expression

Before we can begin quantifying reporter expression in our samples, we need to separate debris from viable cells and then identify the subset of viable cells that are successfully transformed. Given that predominantly healthy cells will express the encoded fluorescent proteins, this property can be used to identify the live cell population (Nadel 1989). Here, we use the RFP signal encoded by the pBeaconRFP vector (Bargmann and Birnbaum 2009) to set an ‘RFP-high gate’ by identifying viable (RFP-positive) cells in a representative transfected sample and comparing it to a non-transformed control (Figure 3). This is accomplished by setting an RFP signal threshold that excludes 99.5% of events in the non-transformed control, using the gate_quantile function from the openCyto package. Events that fall above this threshold in transfected samples are considered RFP-positive. However, not all these fluorescent events represent intact cells as damaged cells or debris may also fluoresce.

**Figure 3:**
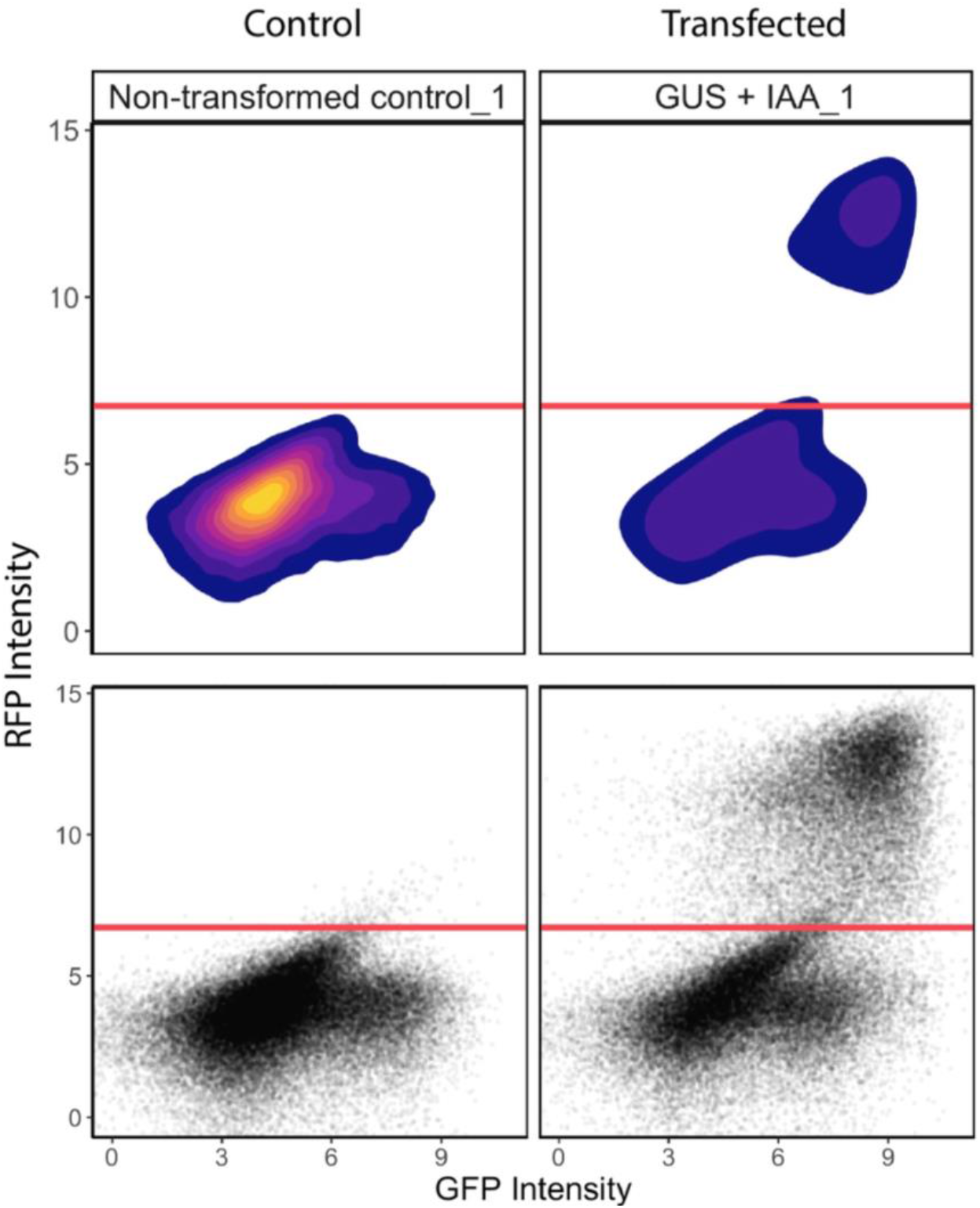
Defining a gate for RFP-positive events. Log scale GFP area signal intensity by RFP area signal intensity depicted as density **(top)** and dot plots **(bottom)** of all events (arbitrary units). Non-transformed control sample **(left)** relative to a representative transfected sample **(right)**. Horizontal red line indicates defined RFP-positive threshold defined as above 99.5% of events in the non-transformed control. Lighter colors represent higher density, and darker colors represent lower density.

To identify the true viable cell population, we back-gate the RFP-positive events onto the FSC-A vs SSC-A distribution generated previously (Staats et al. 2019). Within this distribution, viable cells cluster in a dense group of similar size and granularity (Figure 4). In contrast, debris and abnormal events are more scattered. By establishing a gate that encompasses only the densest 90% of RFP-positive events, we exclude additional debris to retain only viable cells. We define this gate using the gate_flowclust_2D function from the openCyto package. This function implements a robust clustering algorithm, based in the flowClust package, to define a population of cells and then defines an ellipse centered about that population and encompassing a certain quantile of the population. This gate can then be used to enrich for viable cells and exclude much of the dead cells, debris, and bubbles. This back-gating strategy hence also identifies untransformed cells useful for subsequent calculations of transformation efficiency.

**Figure 4:**
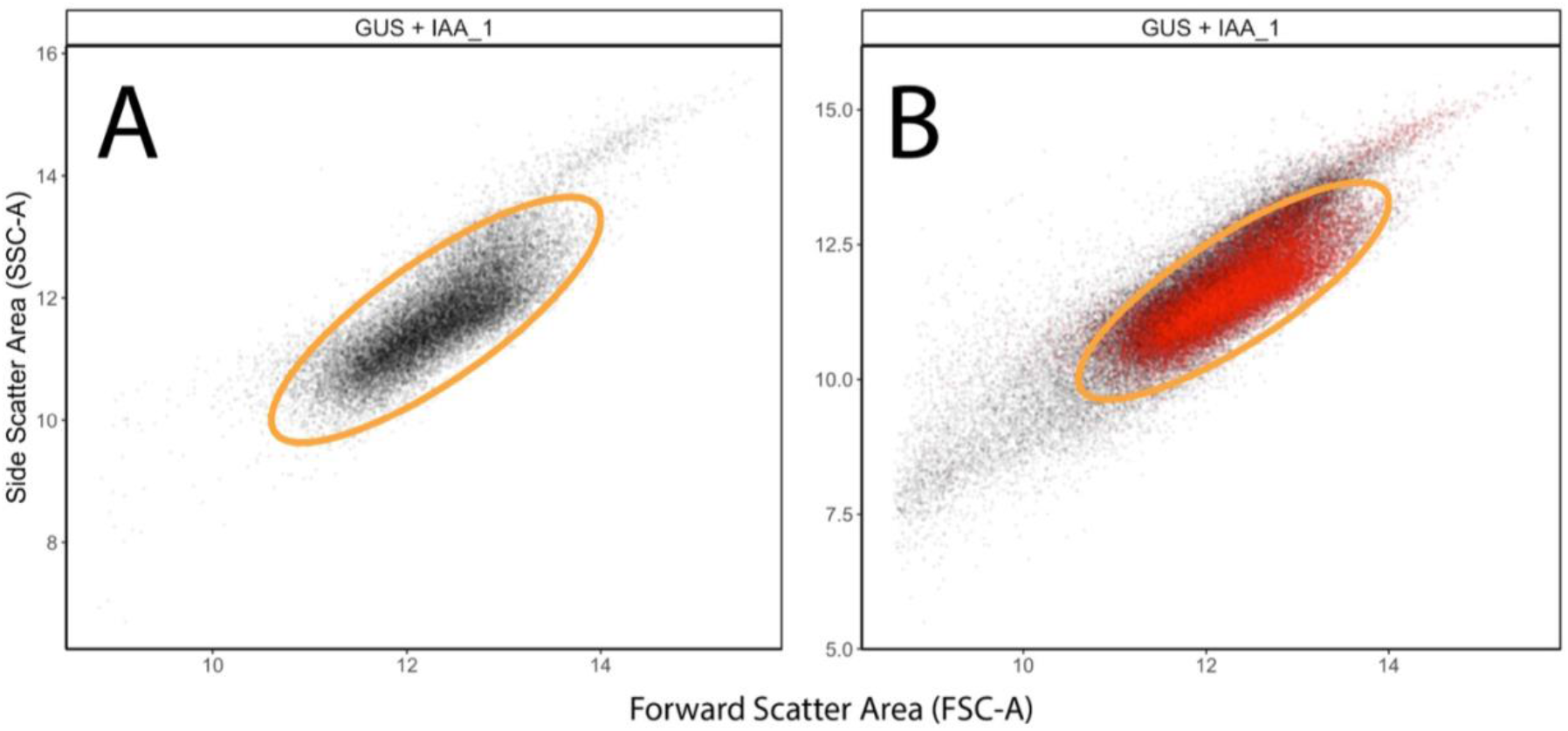
Defining a live-cell gate based on positive RFP expression. **(A)** Log scale forward scatter area by side scatter area dot plot of RFP-positive events. Colored ellipse encompasses 90% of the RFP-positive events, which we define as the viable cells gate. **(B)** Log scale forward scatter area by side scatter area distribution of RFP-positive events plotted against all events. Back-gated RFP-high events (red) overlayed on top all events (black). The viable cells gate encompasses the densest region of all events, as expected in a relatively clean sample.

### Doublet discrimination

Cell doublets (cells that are stuck together) are another factor that may bias our analysis by inflating reporter-gene measurements. Given that protoplasts lack a rigid cell wall, the plasma membrane becomes the outermost boundary, and the cell adopts a near-spherical shape. Consequently, protoplast singlets should exhibit a tight 1:1 correlation between FSC-A (the total signal over time), and FSC-H (the peak signal intensity) (Stadinski and Huseby 2020). In contrast, when two or more cells pass through the cytometer laser in close succession, they generate a broader signal pulse, increasing FSC-A while having less effect on FSC-H. To screen out cell doublets, we select the densest 90% of events in the viable cell population that fall along the expected FSC-A vs FSC-H diagonal (Figure 5) using the gate_singlet function from the flowStats package. With large cell populations, it is generally better to set tighter constraints to minimize the inflationary effect of cell doublets on reporter gene measurements. The 90% threshold is chosen to exclude as many potential doublets as possible while retaining the upper portion of the distribution, which are unlikely to represent doublets.

**Figure 5:**
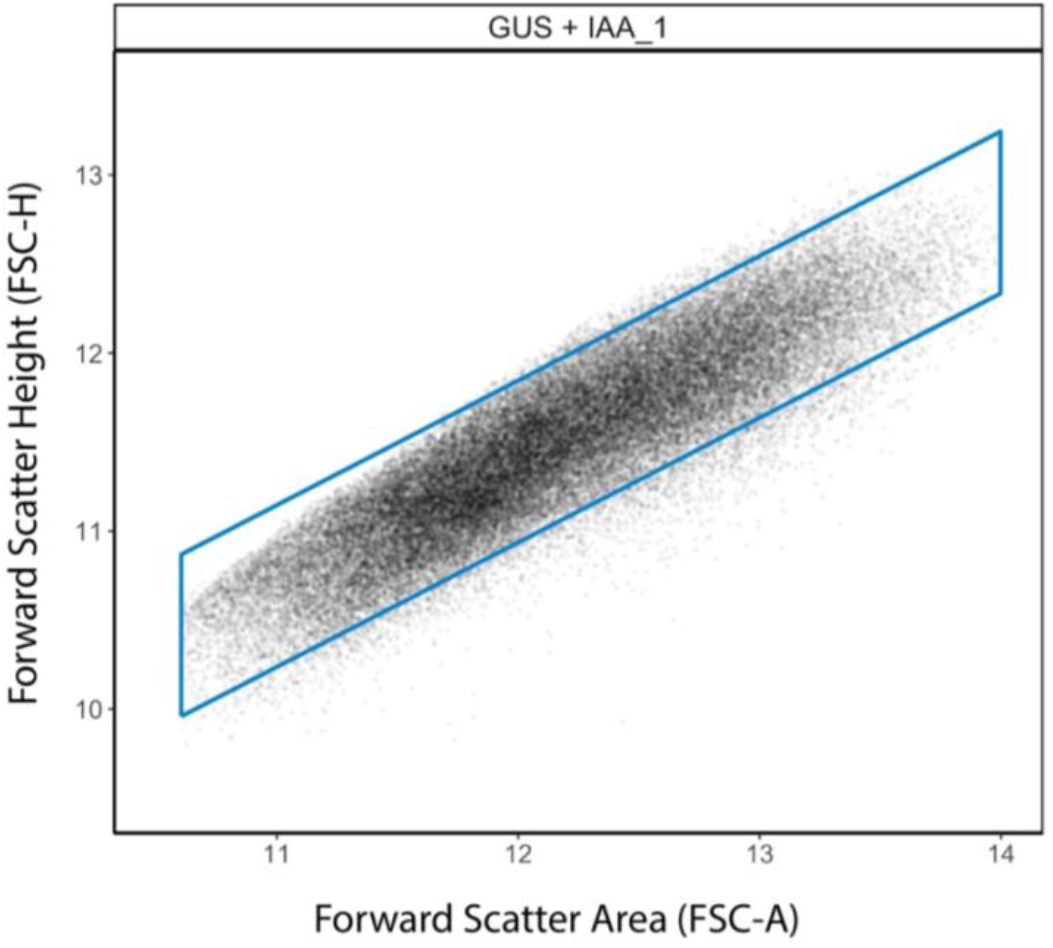
Defining a singlet gate. Log scale FSC-A by FSC-H dot plot of viable cells. The parallelogram (blue) encompasses the densest 90% of cells that follow the FSC-A vs FSC-H diagonal.

### Gating viable, RFP-positive cells

With viable singlet cells isolated, we use this population from the non-transformed control sample to define a refined RFP-positive gate to identify true transformed cells (Figure 6). Given that debris with high autofluorescence has been excluded, the previous RFP-positive threshold is lowered capturing more of the transformed cell population. With the identified population of viable cells that express our effector gene construct, we can now address the initial research question, “How does overexpression of the transfected effector genes affect the genome-integrated GFP auxin reporter activity?”

**Figure 6:**
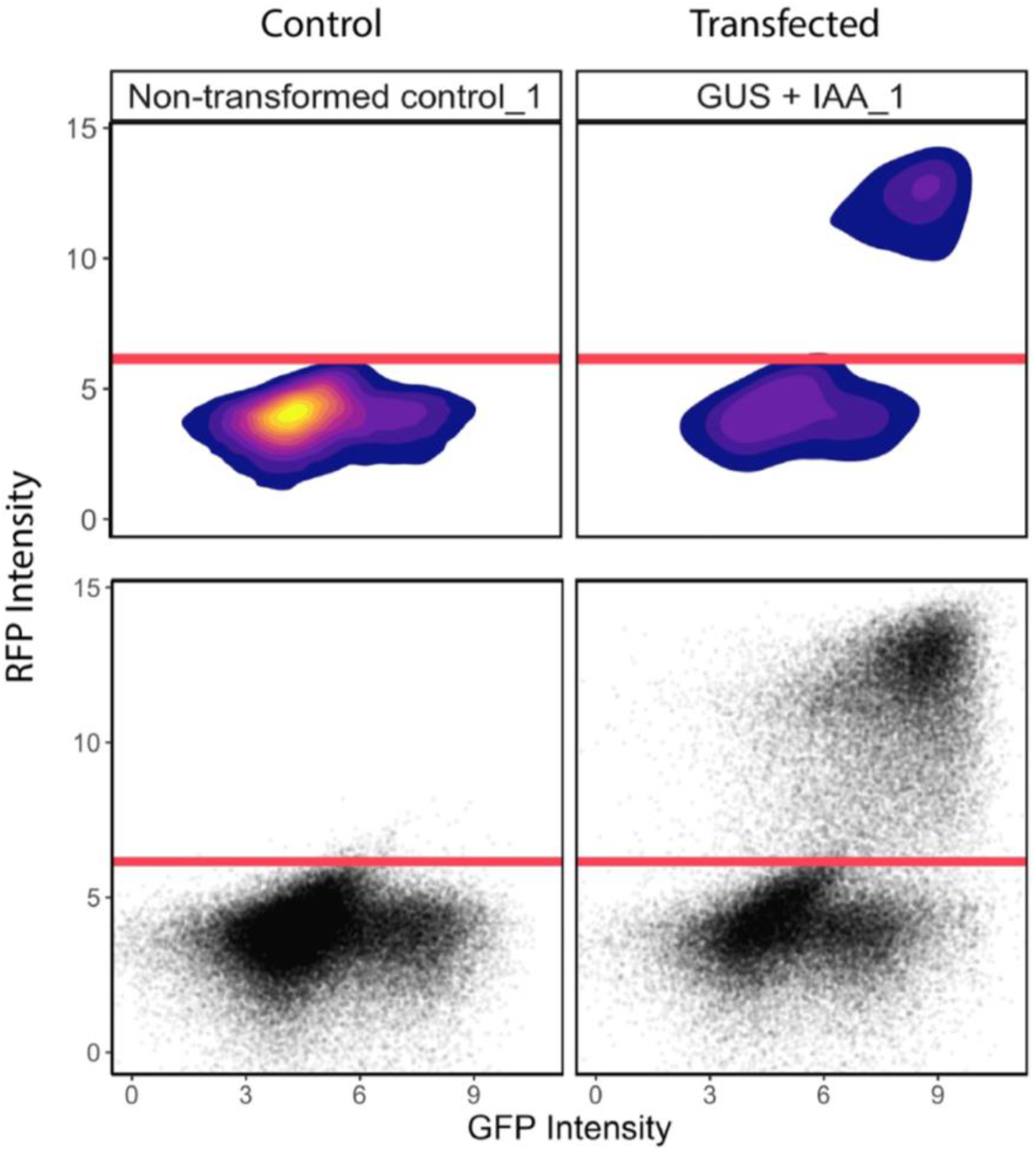
Refining the RFP-positive gate based on the viable singlet cell population. Log scale GFP signal intensity by RFP signal intensity depicted as density **(top)** and dot plots **(bottom)** of viable singlet cells (arbitrary units). Non-transformed control sample **(left)** relative to a representative transfected sample **(right)**. Horizontal red line indicates defined RFP-positive threshold defined as above 99.5% of events in the non-transformed control. Lighter colors represent higher density, and darker colors represent lower density.

### Visualizing replicate distributions and calculating transformation efficiency

With the cell population-of-interest selected, we can verify there are no major differences between sample replicates or transformation efficiency between samples. Sample distributions can be visualized using a density ridges plot of GFP reporter activity (Figure 7A). Given that our samples are technical replicates, the distributions of individual replicates are relatively similar. Dividing transformed cells by the total population of live cells enables us to calculate transformation efficiency (Figure 7B). In our example flowSet, transformation efficiencies range from 26 to 39% while the non-transformed control is 0.5% as expected.

**Figure 7:**
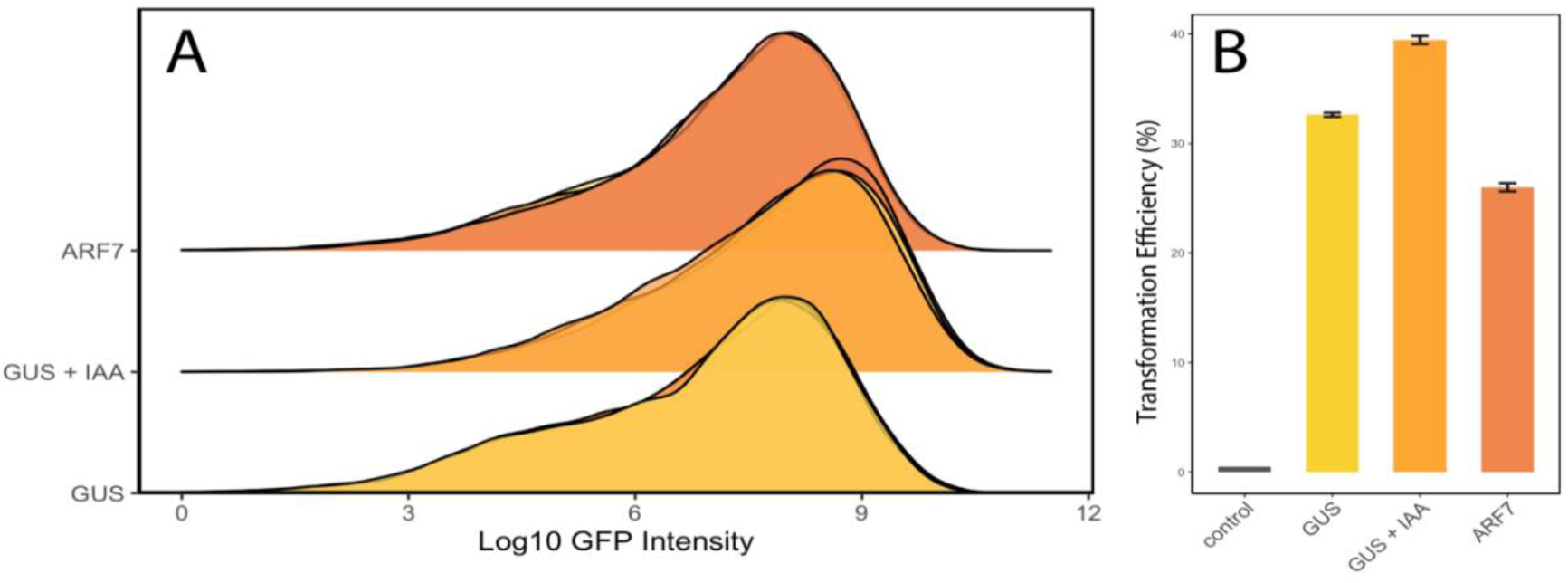
Visualizing replicate distributions and calculating transformation efficiency. **(A)** Log scale density ridges distribution of RFP high cells for each sample with individual replicates overlayed. X-axis is *GFP* activation (arbitrary units). (**B)** Average transformation efficiency (%) by sample. Error bars represent standard error of the mean of three replicates.

### Calculating summary statistics

With replicate consistency verified, we can calculate summary statistics of the sample means using a one-way ANOVA. To run an ANOVA three assumptions must be met: 1) datapoints must be independent from one another; 2) residuals are normally distributed; and 3) homogeneity of variance (Bittner 2022). Independent observation is generally built into the experimental design but can be checked with the Durbin-Watson test for the autocorrelation of observations (Fox 2016). A Durbin-Watson test value of approximately 2 indicates no autocorrelation, while values less than or greater than 2 indicate a positive or negative autocorrelation, respectively. Generally, a Durbin-Watson test is not necessary for randomly ordered biological replicates as independence is assumed by proper experimental design. Importantly, the use of technical replication in our provided vignette technically violates this first assumption and is thus meant to serve only as a simulated example.

Residual normality, referring to differences between observed and group means can be checked with a Q-Q plot. The Q-Q plot is used to compare quantiles of your observed data to quantiles of a theoretical normal distribution. Data that is normally distributed should fall roughly along a straight 45º line (Figure 8) (Wilk and Gnanadesikan 1968; Kozak and Piepho 2017).

**Figure 8:**
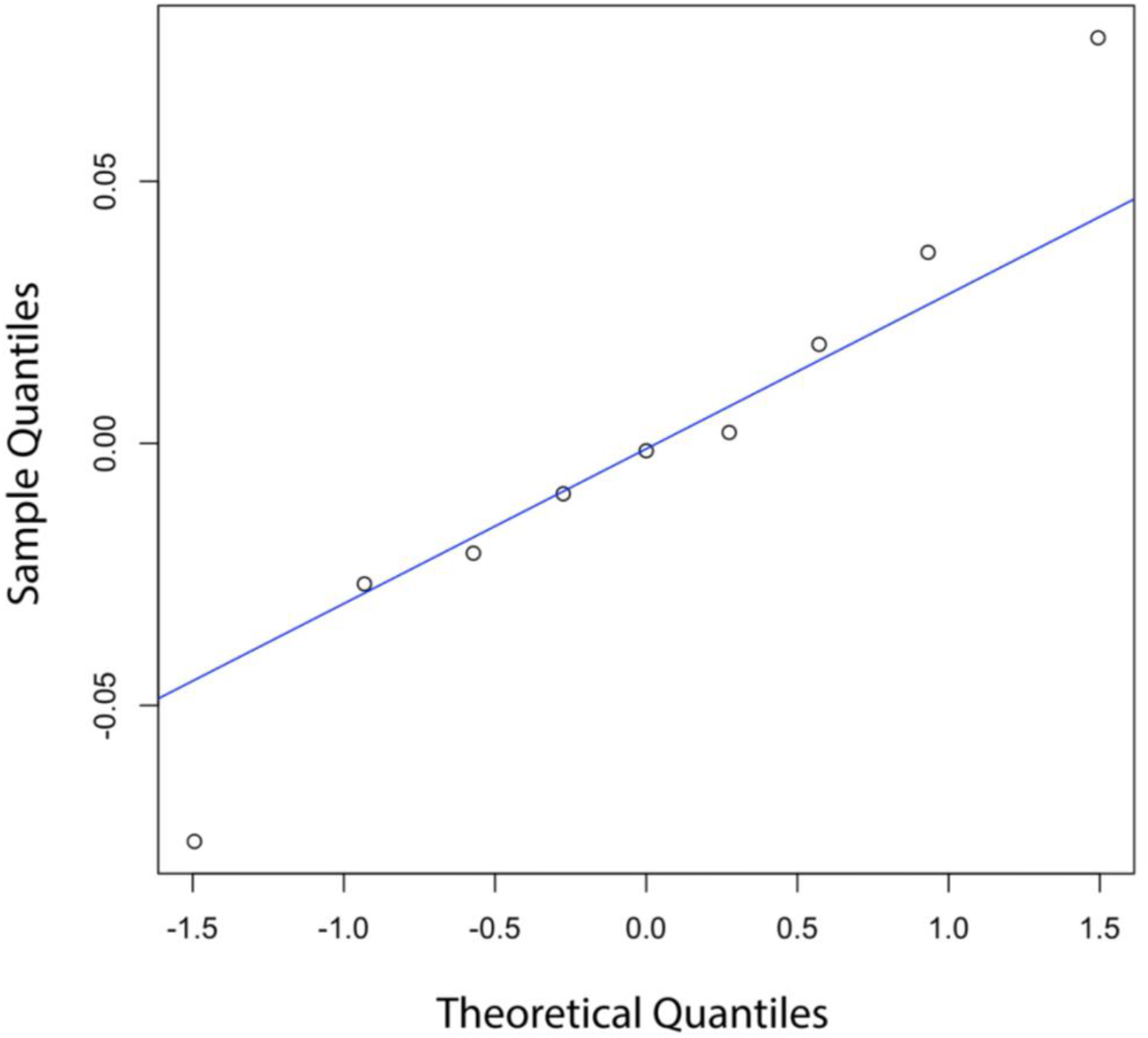
Q-Q plot of model residuals to assess normality. The residuals from the one-way ANOVA model are plotted against the theoretical quantiles of a normal distribution. Points falling near the reference line (blue) indicate that the residuals are approximately normally distributed, supporting the assumption of normality required for ANOVA.

The final assumption, homogeneity of variances, assumes the variance within each group should be roughly equal. Variance can be evaluated using Levene’s test which determines whether the variance in the dependent variable (e.g., reporter gene activation) is roughly the same across samples (Levene 1960). A Levene’s test returning a p-value > 0.05 indicates that the assumption of equal variances is met. If all the assumptions are met, we can use the one-way ANOVA with a Tukey post-hoc HSD test to compute pairwise comparisons between samples.

If not all the assumptions of an ANOVA are met, some options are available depending on which assumption is violated. Generally, independence of observation should be ideally addressed by experimental design, but if the data is hierarchical or nested, it can be accounted for by fitting the data to a mixed-effects model (e.g. the lme4 package in R). If the data is not normally distributed, data transformations (e.g. log, square root, or Box-cox) can help improve normality. Alternatively, non-parametric tests like the Kruskal-Wallis test provide an alternative option that does not assume normality. If the equal variances assumption is violated, Welch’s ANOVA is a viable alternative as it adjusts for unequal variances.

### Quantifying reporter-gene activation and generating figures

Violin and boxplots are powerful visualization tools that can be overlaid to display the distribution, median, mean, and outliers of each sample in a single figure (Figure 9A). In this case, we can plot log-transformed GFP intensity values to reduce skew, stabilize variance across samples, and align the visualization with the scale used for statistical testing. However, log transformation also compresses the visual scale, particularly at higher values, making statistically significant differences appear less striking (West 2022). To improve interpretability, we can back-transform the estimated means and calculate fold-change in *GFP* activation relative to the inert control (*GUS*) as a baseline (Figure 9B). Given that differences on a log scale correspond to fold-changes on a linear scale, we can retain the statistics calculated on the log-transformed data while presenting more visually interpretable differences.

**Figure 9:**
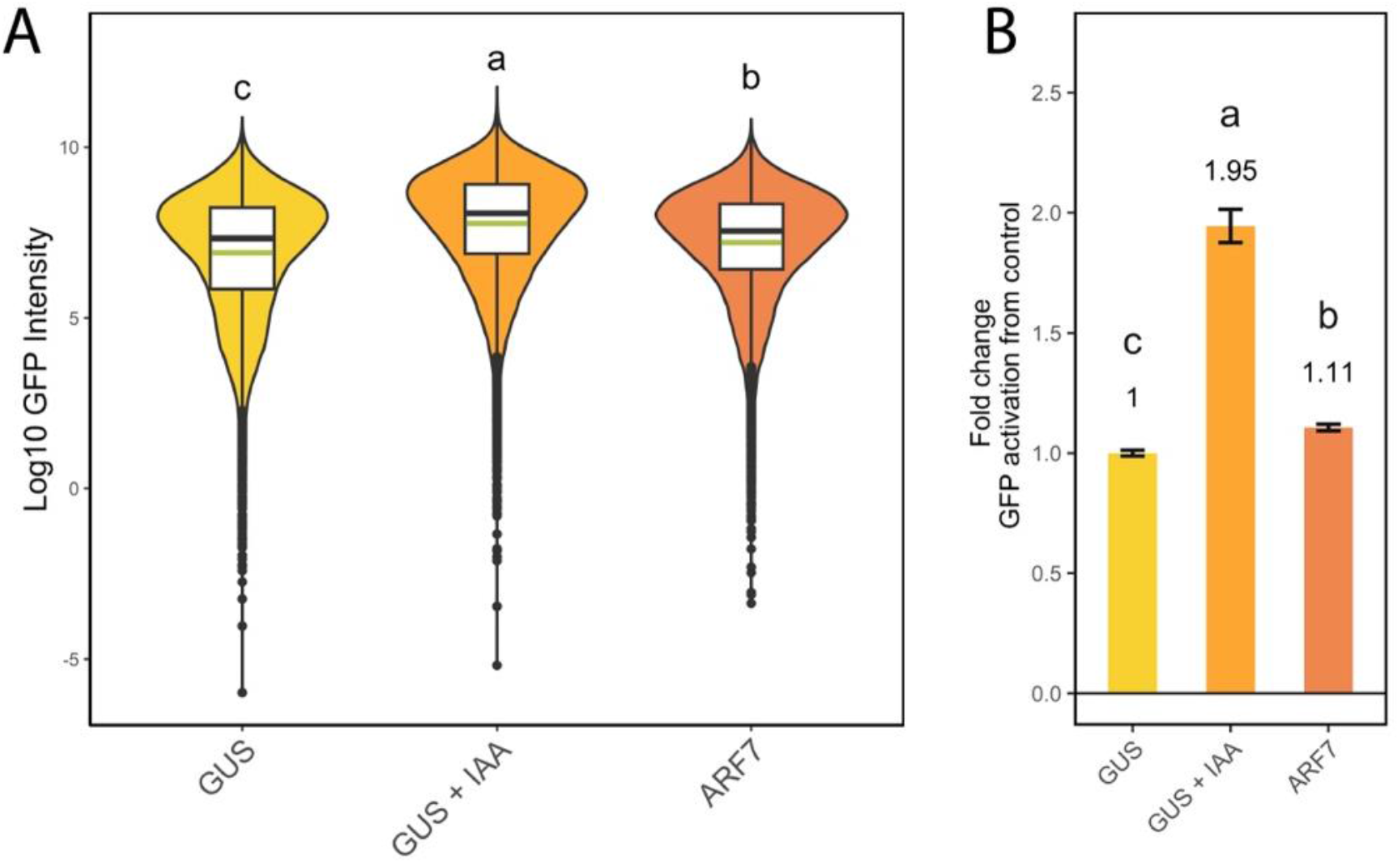
Comparing reporter-gene expression between transformation and treatment conditions. **(A)** Box violin plot of Log scale *GFP* activation of transformed (RFP-positive) cells. Median is shown in black. Mean is shown in green. **(B)** Back-transformed average fold change in *GFP* activation relative to *GUS* control sample in RFP high cells. All statistical tests were performed on log_10_-transformed fluorescence intensities, which stabilized variances across groups. Letters indicate groupings based on Tukey-adjusted pairwise comparisons; constructs sharing a letter are significantly similar (One-way ANOVA, α = 0.05, Levene’s *p* > 0.05).

## Results and Discussion

In the worked example, we observed the influence of different effectors on the auxin reporter, *DR5::GFP*. While the dataset’s primary purpose is to serve as an example of data processing, the results also align with other protoplast studies involving auxin signaling, validating the workflow’s utility. Each sample exhibited statistically significant differences which were nearly imperceptible at log-scale due to the data spread (Figure 9A). When plotted relative to the inert control (*GUS*), we observed that GFP reporter activity was elevated in both the auxin-treated and *ARF7/NP4* samples which lead to 1.95-fold and 1.11-fold statistically significant increases respectively (Figure 9B). This aligns with the findings of Tiwari et al. (2003) who demonstrated that *ARF7/NPH4* encodes a transcriptional activator that is partially repressed by Aux/IAA proteins in the absence of auxin. This is consistent with our data in which overexpressed *ARF7/NPH4* exhibits reduced reporter-gene transcription relative to the auxin treatment.

Interestingly, *ARF7/NPH4* and more prominently *GUS* have slightly bimodal distributions that is not present in the auxin-treated sample (Figure 7A and 9A). This may be explained by the heterogenous mixture of root cells that make up the dataset. Thus, subpopulation-specific regulatory circuits within these cells may enhance or suppress the reporter gene activity, creating separate high and low *DR5::GFP* expressing populations. Auxin-treatment may override such circuits, explaining why we do not see a bimodal distribution in the *GUS* + IAA sample.

While this workflow improves the reproducibility and scalability of transient protoplast expression assays, several limitations should be considered. First, the workflow does not include steps for calculating the fluorescence compensation matrix. Instead, it assumes that compensation settings have already been calculated by the cytometer’s acquisition software and applies them. While this is common practice, it introduces potential variability and reduces transparency in how spectral overlap between fluorophores is corrected. Future iterations of the workflow could integrate automated or user-guided compensation tools directly within the R environment to support fully end-to-end analysis.

Second, while the use of dual fluorescence and numerically defined gating improves objectivity and reproducibility, the accuracy of gating decisions can still be influenced by user-defined thresholds or parameters. Although we provide visualization tools for quality control, some manual tuning may still be required in challenging datasets (e.g., low transformation efficiency or overlapping populations).

Third, although the workflow streamlines many aspects of flow cytometry data analysis, it is not fully automated. Users are still required to set key parameters, visually inspect gating results, and manually adjust code for dataset-specific features. While this design encourages transparency and flexibility, it may limit applications where minimal user intervention is desired. Future iterations of this workflow could improve automation for batch processing and standardized report generation.

## Conclusion

We have assembled a workflow to process transient-expression flow-cytometry data using the R programming language and core flow-cytometry infrastructure of the Bioconductor R package repository. This workflow allows researchers with basic levels of cytometry and programming experience to: 1) annotate, compensate, and transform the dataset, 2) distinguish viable cells from bubbles and debris, 3) identify successfully transformed (RFP-positive) cells, 4) measure integrated GFP reporter activation in transformed cells, 5) conduct statistical analysis, and 6) generate publication-ready figures. Critically, this workflow allows for a fully transparent, open-source, system-agnostic, high-throughput, and reproducible data analysis. The workflow is available at: https://github.com/PlantSynBioLab/positive-fluorescence-selection.

## Acknowledgements

We are grateful to Emilia Hyland and Virginia Tech’s Center for Biostatistics and Health Data Science (CBHDS) for their helpful feedback. Research in the Bargmann lab is supported by U.S. National Science Foundation (United States) IOS 2343702, NSF IOS 2314549, USDA National Institute of Food and Agriculture, HATCH project VA-160260, and Multistate S-009 project VA-136423. Research in the Wright lab is supported by U.S. National Science Foundation (United States) IOS 2343702, United States Department of Agriculture National Institute of Food and Agriculture (USDA-NIFA) Hatch Project VA-1021738, and the National Institute of General Medical Sciences of the National Institutes of Health under award number R35GM150856. The content is solely the responsibility of the authors and does not necessarily represent the official views of the National Institutes of Health. E.A.S. received funding from the Fralin Undergraduate Research Fellowship program (Virginia Tech). J.S.T. is supported by the George Washington Carver Program (Virginia Tech).

