## Supplementary material for "Glow with the Flow: Reproducible Analysis of Transiently Transformed Protoplasts Using Dual Fluorescent Reporters in R": Protoplast_Automation.Rmd

````
---
title: "A Standardized Workflow for Flow Cytometric Data Analysis of Protoplast Transient Expression Assays"
author: "Joseph Taylor, Elyse Shoppell, Clay Wright, Bastiaan Bargmann"
date: "4/24/2025"
output:
  html_document: default
  pdf_document: default
---
## Abstract

Transient transformation assays using protoplasts have become a widely employed technique in plant research. Positive fluorescent selection of successfully transformed cells was subsequently developed to assess the effect of transient effector gene expression in only successfully transfected cells using flow cytometry. This process, though effective, often requires considerable manual effort and subjective judgment to quantify reporter gene expression in the intended cell populations. To address this, we introduce a new, open-source workflow based on the R programming language. This method enhances the reproducibility and scalability of such experiments, which enable rapid study of gene regulation and signal transduction in plants.

## Introduction

Transient transformation of protoplasts is a widely used technique in plant research due to its relative ease and utility (Bargmann and Birnbaum 2009; Nanjareddy et al. 2016; Lin et al. 2018). An advantage of this technique is the speed and throughput with which experiments can be conducted in addition to enabling quantitative measurements of reporter-gene activation. These assays are often used to quantify signaling and gene regulatory networks by assessing gene expression, promoter elements, transcription factor or signal-transduction protein activity, and subcellular protein localization (Tiwari et al. 2006). Although transient assays in protoplasts to assess reporter gene activation have proved invaluable in elucidating the elements involved in the regulation of gene expression, the approach comes with limitations. Historically, transcription factor studies using protoplasts involved co-transfection with up to three plasmids: 1) a promoter::reporter construct (e.g., β-GLUCURONIDASE or firefly LUCIFERASE); 2) an effector construct (e.g., a transcriptional activator/repressor); and 3) a constitutively expressed transformation reporter (e.g., Rinella LUCIFERASE/RLuc) (Ulmasov et al. 1997a;b). A drawback of these initial systems was the potential for skewed measurements due to variations in protoplast transformation efficiency (Liu et al. 1994). These assays were later adapted to use a genome-integrated reporter system, forgoing the need for triple co-transfection, reducing experiment complexity. However, the field still lacked the ability to selectively measure only successfully  transformed cells, which typically represent a subset of the total population (Tiwari et al. 2006).

Positive fluorescent selection of protoplasts was developed as a contemporary approach to quantify transient effector gene expression in only successfully transformed cells via flow cytometry (Bargman and Birnbaum 2009). This system allows for analysis of only the transformed cell population, requiring only a single vector input coding for the effector (pBeaconRFP/GFP).  In addition to the effector, the vector also encodes a fluorescent protein reporter used for identification of transformed cells (e.g., red fluorescent protein, RFP), as well as the effector gene. The activation of a genome-integrated fluorescent reporter (e.g., green fluorescent protein, GFP) is then measured in only those RFP-positive cells expressing the transfected effector gene. Furthermore, the addition of fluorescence-activated cell sorting of the successfully transformed cell population enables additional downstream analyses (e.g., transcriptome analysis via RNA sequencing). 

Flow cytometry paired with positive fluorescent selection enables rapid, single-cell analysis of transient expression in protoplasts (Bargman and Birnbaum 2009; Alverez et al. 2020; Beard et al 2021; Gonzalez et al. 2021). However, reproducibility of data analysis is crucial (Sandve et al. 2013). In typical implementations, proprietary flow cytometry analysis software is used to identify the transformed-cell population and quantify reporter gene expression (Sony Corporation 2015; ThermoFisher Scientific 2015). Additionally, the identification of viable, successfully transformed cells using standard flow cytometry software is time-consuming and relies heavily on subjective gating decisions (Bio-Rad 2023). As a result, we developed an open-source, data-driven analysis workflow using the R programming language to improve the reproducibility and scalability of transient expression assays for investigating reporter gene activation. Here, we present example vignette using this workflow to standardize and semi-automate analyses for high-throughput (e.g., 96-well plate) transient expression experiments using dual-fluorescent reporter systems (e.g. GFP and RFP). 

We have assembled this workflow using the core flow cytometry infrastructure of the Bioconductor R package and have also deposited the workflow and vignette in this repository (Hahne et al. 2009; 2025; Finak et al. 2014; Wright et al. 2025; Van et al. 2015). While R programming requires the use of a text/command-line-based interface, it also offers the ability to develop templated reports with integrated descriptive text, R code, and figures. This format, known as R-markdown, provides an ideal medium for transparently communicating the complete analysis and results in a single, easily reproducible format (Allaire et al. 2023; Xie et al. 2023) Additionally, we have found the community of R users to be very welcoming and helpful to new learners, as evidenced by the numerous, free, community-developed resources for learning (find a short list of excellent resources at https://education.rstudio.com/learn/beginner/). This workflow can be broken down into a series of steps: 1) Annotate, compensate, and transform the dataset, 2) distinguish individual cells from bubbles and debris, 3) identify successfully transformed (RFP-positive) cells, 4) measure reporter activation (DR5::GFP) in transformed cells, 5) conduct statistical analysis, and 6) generate figures. The included vignette uses the R-markdown format where each step of data processing and analysis is described in greater detail. For much of this workflow, we utilize the flowCore, flowStats, and flowTime packages from Bioconductor. To define data-driven gates we use a clustering algorithm, as implemented in the Bioconductor packages and openCyto [@Finak2014]. This allows us to define rectangular or elliptical gates which contain a certain quantile of a specified population. These algorithmically defined gates are then plotted for visual inspection and potential adjustment of the quantile parameter.

Ultimately, this workflow enables a fully transparent, open source, and reproducible data analysis pipeline for transient expression assays, adaptable to various flow cytometry platforms. Annotation steps included in the workflow facilitate attachment of experimental metadata to the dataset according to FAIR principles [@wilkinson_fair_2016] as required by FlowRepository [@spidlen_flowrepository_2012], a central repository of flow cytometry datasets. We have validated this workflow with the included example dataset and demonstrated its utility in a peer-reviewed publication (Taylor et al. 2025). While here we focus on experiments using plant protoplasts, this analysis is applicable to any cell type including yeast, mammalian cells, and fungal protoplasts.

## Methods


### Example dataset overview
The worked example data consists of Arabidopsis root protoplasts carrying an integrated green-fluorescent auxin reporter (DR5rev::GFP). These cells were transfected with the pBeaconRFP vector, which encodes RFP and an effector gene–either AUXIN RESPONSE FACTOR 7/NON-PHOTOTROPIC HYPOCOTYL 4 (ARF7/NPH4) or β-GLUCURONIDASE (GUS, an inert control)–or were mock-transformed. Additionally, a subset of the cells transfected with the GUS construct were also treated with 100 nM indole-3-acetic acid (auxin). 

### Transient expression assay and flow cytometry
Described similarly in Gonzalez et al. (2021) and Bargmann and Birnbaum (2009), roots of one-week-old seedlings were harvested and placed into a 500 mL flask, gently shaking at approximately 75 RPM and 28 ºC in 50 mL enzymolysis solution (1.25% [w/v] Cellulase R-10 [Yakult Pharmaceutical Industry Co., Japan], 0.3% [w/v] Macerozyme R-10 [Yakult Pharmaceutical Industry Co., Somerset, NJ, USA], 0.4 M mannitol, 20 mM MES, 20 mM KCl, 10 mM CaCl2, 0.1% [w/v] bovine serum albumin; pH was adjusted to 5.7 with KOH pH 7.5) for 5 h. The protoplast solution was filtered through a 40 µm Falcon cell strainer (VWR, Radnor, PA, USA), divided over 15 mL conical tubes, and centrifuged for 5 min at 500 g. Cells were gently washed in resuspended in the enzymolysis buffer (no enzymes added), centrifuged for 5 min at 500 g, resuspended in 1 mL of the enzymolysis buffer, and then counted with a hemocytometer. Cells were then washed with transfection solution (0.4 M mannitol, 15 mM MgCl2 hexahydrate, 4 mM MES; pH was adjusted to 5.7 with KOH), centrifuged again, and resuspended in transfection solution with a final density of 4x106 protoplasts/mL. Conical tubes (15 mL) were prepared for each transfection with 25 µg of plasmid DNA (1 µg/µL) and 250 µL of protoplasts in transfection solution. Protoplasts were transformed with the pBeaconRFP vector carrying ARF7/NPH4, GUS, or not transfected. Next, a 250 µL PEG solution (40% [w/v] PEG 1500, 0.2 M mannitol, 0.1 M CaCl2) was added, and the suspension was mixed by flicking the tube repeatedly. Suspensions were immediately washed with 15 mL of enzymolysis buffer, centrifuged, and resuspended in 750 µL enzymolysis buffer. One of the two GUS samples was treated with 100 nM indole-3-acetic acid. Protoplast suspensions were then divided three ways in a 96-well plate (250 µL per well) to generate three technical replicates. The plate was incubated overnight in the dark at room temperature. 

The GFP intensity in RFP-positive cells was quantified using a Sony SH800S fluorescence activated cell sorter with 488 nm and 561 nm lasers measuring fluorescent emission at 525/50 and 617/30 nm for GFP and RFP/mCherry fluorescence, respectively. Data was exported from the proprietary cytometry software in the flow cytometry standard file format (.FCS). The R markdown was developed and tested using R version 4.4.3. 

## Getting Started

Export the flow cytometry dataset as .fcs files from your flow cytometer. To set up your work space, create a directory (folder) with a name of your choosing. In this parent folder, place this R markdown document, and a create a sub folder titled "FCS_files" where you should deposit all of your .fcs files. If this script is not in the same parent folder as the FCS_files folder, you will need to set the working directory to tell R where your files are. Use `setwd(" ")` with the file path to the parent folder folder (see example below) to set the working directory. If your experiment spans across multiple multi-well plates, some of your samples may have identical file names, you should separate them with additional sub folders (i.e. 'plate A', 'plate B') or manually change the file names.  

First, we will install required the packages.  

```{r}
### ------------------  Installing Required Packages ------------------ ###


# Use the following lines of code to install necessary packages (This may take approximately 5 minutes, but you should only need to install the packages once):

# install.packages(c("tidyverse", "ggbeeswarm", "dplyr", "forcats", "ggridges", "agricolae", "hrbrthemes", "viridis" ,    "BiocManager", "htmltools", "cli", "fastmap", "deSolve", "Hmisc", "car"))

# All of the packages below are in the Bioconductor repository
# BiocManager::install(c('flowStats','openCyto','flowTime','ggcyto','flowCore'))

# If you are prompted to update any of the packages, please do so.

```

Now that each required package is installed we need to load it for the current R session. 

```{r results = "hide", message = FALSE, warning = FALSE}
### ------------------  Loading Required Packages ------------------ ###

# List of packages
packages <- c(
  "flowTime", "flowCore", "flowStats", "openCyto", "ggcyto", 
  "tidyverse", "ggbeeswarm", "dplyr", "forcats", "ggridges", 
  "htmltools", "agricolae", "hrbrthemes", "viridis", "car"
)


# Load all packages
lapply(packages, require, character.only = TRUE)


```

Once your work space is set up, we need to tell R where our files are on the computer. We can do this by defining the working directory using `setwd()` (e.g. `setwd("~/TransientExpression/2022/ExampleExperimentFolder")`). 

Next, we will define an `experiment_name` and `experiment_date` to be assigned to our flowset once it has been annotated.

```{r}
### ------------------  Setting the Experiment Name and Date  ------------------ ###


# Set the experiment name and date
experiment_name <- "Example_Data"  # Experiment Name

experiment_date <- "10-16-24" # Experiment Date

```

## Import the raw data

Now we can import the raw .fcs files as a `flowSet` object into the R environment. If you have a large dataset, this may take a minute or two.  

```{r}
### ------------------  Importing Raw FCS Data ------------------ ###


# Read in the flow cytometry dataset and assign it to the object "fs".
fs <- read.plateSet(pattern = "FCS_files", truncate_max_range = FALSE) 

```

## Annotate the dataset

Importing the FCS data as a flowSet into the R environment using the provided workflow, allows for annotating, compensating, and converting the flow cytometry files into a format which can be used to generate publication-ready figures and calculate summary statistics.
Annotations are important for enabling repeatable analyses by independent researchers. If the annotations meet or exceeds the minimal information standards for flow cytometry experiments defined by [@lee_miflowcyt_2008], then the dataset can be deposited into FlowRepository [@spidlen_flowrepository_2012] to be shared with others in the scientific community. Annotations can include information such as sample name, expression vector, treatment type, duration, concentration, etc. Annotated flowSets can be exported, allowing others to reload the dataset with experiment information preserved for future use.


First, we will create a data frame as a template with columns "name" (the .fcs file name), "construct", "treatment", and "replicate," and any other parameters you wish to include before exporting it as a .csv file. Using excel or another spreadsheet editor, you should then fill in any information for each sample you would like included in the annotation. If you do not have any information for a certain variable (i.e. treatment), you may leave it blank or delete that column entirely. All sample "names" in the .csv should exactly match the .fcs file name (i.e. 'A7-1.fcs') for the annotation to work properly. Once you have finished filling out your .csv file, we can then apply the annotation to the flowSet `fs` as follows:   

```{r}

### ------------------  Defining Sample Names and Treatments ------------------ ###

# # Create a template data frame for annotations
# annotation_template <- data.frame(
#   name = NA,
#   construct = NA,
#   treatment = NA,
#   replicate = NA
# ) [numeric(0), ]
# 
# # Export the empty template data frame to a .csv file
# write.csv(annotation_template, file = "sample_info.csv", row.names = FALSE)

# Read in the annotated .csv file 
annotation <- read.csv("sample_info.csv") 

# Note: You may receive the following warning if the last line of the .csv file is not blank: "Warning: incomplete final line found by readTableHeader on 'sample_info.csv'." This is not an error and can be ignored, the .csv has still been properly imported into R.  

# Note: FlowTime adds a leading "1" to the name of each .fcs file, so we will to adjust our annotations to match
annotation <- annotation %>% mutate(name = paste0("1", name))

# Apply the annotation the to dataset
fs <- annotateFlowSet(yourFlowSet = fs, annotation_df = annotation, mergeBy = 'name')
```

We can also add additional variable metadata, which allows us to add more detailed descriptions for each variable in your annotation, to the annotated flowSet. For example, in addition to the experimental treatment variable, you may want to include specific details about each treatment such as duration, concentration, source, etc. 

```{r}
### ------------------  Add Additional Variable Metadata ------------------ ###

# Add additional metadata to each variable and apply it to the annotated flowset. 
vMd <- varMetadata(fs@phenoData) 
vMd['construct','labelDescription'] <- "Example: All constructs are driven by the 35S promoter in the pBeaconRFP vector"
vMd['replicate','labelDescription'] <- "Example: Each sample has three biological replicate "
vMd['treatment','labelDescription'] <- "Example: Auxin-treated samples were treated with 100 nM indole-3-acetic acid (IAA/auxin) overnight"

# Apply these annotations to the flowset
varMetadata(fs@phenoData) <- vMd

```

## Save the annotated dataset

Now we can save this annotated flowset which will allow anyone in the future to load it in with all of the annotations. The flowset is exported as .fcs files and the annotations are saved as 'annotation.txt.' This will all be saved in a new folder based on the name of your experiment and the date within the parent folder that you set up at the beginning of this document. Clearly annotated datasets improves the reusability of datasets and reproducibility of data analyses.
 
```{r, eval=FALSE}

### ------------------  Export the Annotated Dataset ------------------ ###

# Save the annotated dataset 
if(!dir.exists(paste0(experiment_date, "_", experiment_name))) {
  write.flowSet(x = fs, outdir = paste0(experiment_date, "_", experiment_name))}

```

## Read in an Annotated Dataset

When we already have a annotated dataset, we can import it here. We use the `phenoData` argument to specify the annotation data frame you created above ('annotation.txt') containing the sample information and experimental metadata.

```{r}
### ------------------  Import an Already Annotated Flowset ------------------ ###

# # Set the experiment name and date
# experiment_name <- "Example_Data"  # Experiment Name
# 
# experiment_date <- "10-16-24" # Experiment Date


# Load an annotated flowset
fs <- read.flowSet(path = paste0(experiment_date, "_", experiment_name), phenoData = "annotation.txt")

# Adding an 'id' column to our flowset that contains both construct name and replicate number to for plotting
pData(fs)$id <- with(pData(fs),
                     paste0(construct, "_", replicate))
```

Now that our flowset is adequately annotated we can begin visualizing the data. However,many makes and models of cytometers use slightly different keywords to reference laser channels, forward scatter and side scatter detectors. To determine which keywords we need to use, we can print all keywords associated with a .fcs file into the R console using `keyword(fs[[1]])`. 

Here, we have set up a few presets that enable us to quickly swap between different keywords in one code chunk rather than every instance it appears in this script. These presets assign cytometry platform-specific keywords to the following variables: Red Fluorescence Channel = `RedDetector`; Green Fluorescence Channel = `GreenDetector`; Forward Scatter Detector = `FSC_A` (Note: some cytometers may refer to 'side scatter' as 'back scatter'); and  Side Scatter Detector =`SSC_A`. The default presets assigns keywords used by the the Sony SH800S fluorescence-activated cell sorter (1), or the ThermoFisher Attune NxT flow cytometer, can be tailored to support different flow cytometry platforms. We can select a defined preset by assigning a either a "1" or "2" to the `CytometerPreset` variable.

```{r}
### ------------------  Define Relevant Keywords ------------------ ###

# Assign preset to 1 (Sony SH800S) or 2 (ThermoFisher Attune NxT)
CytometerPreset <- 1

# Conditional to set the keywords based on the CytometerPreset value
if (CytometerPreset == 1) {
    # For Sony SH800S fluorescence-activated cell sorter
    RedDetector <- "FL3.A"
    GreenDetector <- "FL1.A"
    FSC_A <- "FSC.A"
    SSC_A <- "SSC.A"
} else if (CytometerPreset == 2) {
    # For ThermoFisher Attune NxT flow cytometer
    RedDetector <- "YL1.A"
    GreenDetector <- "BL1.A"
    FSC_A <- "FSC.A"
    SSC_A <- "SSC.A"
} else {
    # Handle the case where CytometerPreset is not set to either 1 nor 2
    print("Invalid CytometerPreset value. Please define the variables, `RedDetector`, `GreenDetector`, `FSC_A`, `SSC_A` as strings (ex. `RedDetector <- 'FL3.A').")
}

```

## Visualize the Raw Data

Now that we have annotated our dataset and assigned relevant keywords, We will plot all of our samples as a quick quality control step to ensure each samples have similar distributions on a forward scatter area vs. side scatter area plot. For larger datasets, this step make take awhile. To look at individual or a subset of samples we can use a subset operator (e.g. `fs[1]` or `fs[1:3]`). The axes of the plot by changing the values for the `xlim` and `ylim` arguments. 

Images of each plot can be exported as JPEGs using the `jpeg()` function 
Image Export Syntax: 

`jpeg("FILENAME", width=####, height = ####, res = ###)`

-code to generate plot-

`dev.off()`

We initially check event distributions of each sample on a log scale to screen for any inconsistencies, possibly caused by contamination or mechanical issues. This is done by generating forward scatter area (FSC-A; event size) versus side scatter area (SSC-A; internal complexity) dot plots (Figure 1). Log transformation of the data is useful to address skewness to achieve a near-normal distribution. FSC and SSC area is used instead of height because it provides more reliable measures of these parameters. Issues at this step may manifest as inconsistent clusters of events between samples in which case the compromised sample could be removed. Given that all samples in the provided flowSet are from the same source tissue and free of contamination, each sample distribution appears similar (Figure 1). 

```{r message = FALSE, warning = FALSE}
### ------------ Plot all events of each sample (FSC.A v SSC.A) ------------ ### 

# Initializes graphics device for exporting the plot as a JPEG
# jpeg("Sample_Distribution_Check.jpg", width=8000, height=6000, res=600)

# Plots all samples in the flow set on a log transformed FSC.A v SSC.A dot plot
ggcyto(data = fs[], mapping = aes(x = !!FSC_A , y = !!SSC_A)) + 
  # Create a density gradient
  stat_density_2d(aes(fill = after_stat(level)), 
                geom = "polygon")+
  scale_fill_viridis_c(option = "plasma") +
  # Uses the id column for plot header
  facet_wrap(~id) +  
  # Sets X and Y axis limits
  coord_cartesian(xlim = c(5e3, 5e6), ylim = c(5e2, 5e6)) +
  # Log transforms the X and Y axis
  scale_x_log10() + 
  scale_y_log10() +
  # Add X and Y axis labels
  labs(x = "Forward Scatter Area (FSC-A)", y = "Side Scatter Area (SSC-A)") +
  # Define plot graphical style 
  theme_classic() +
  theme(
  panel.border = element_rect(color = "black", fill = NA, size = 1))+
  theme(
    strip.text = element_text(size = 14),      # plot header
    axis.title = element_text(size = 16),      # X and Y axis labels
    axis.text = element_text(size = 12)        # axis tick labels
  )

# Closes the current graphics device. This finalizes and writes the image to disk
# dev.off()   

```


## Compensate the flowSet

The first step in our data analysis is to compensate for any spillover of each fluorescent protein used into channels used to measure another fluorescent protein. For example, it is often the case that RFP is measured in the FL3 (YL1 in some cytometers) channel, but it also contributes some signal in the FL1 (BL1 in some cytometers), where GFP is measured, therefore, cells expressing high levels of RFP will have inflated GFP measurements. In some cases, the emission spectra and measurement channels of the two fluorescent proteins may not result in any spillover. To determine the amount of spillover and calculate a compensation matrix, control samples that express each fluorescent protein individually and also samples with no fluorescent proteins are required. 


Only a small fraction of cells are likely to express the fluorescent protein within a population of transfected cells, so we need to first identify this cell population and determine the spillover in just this population. Some cytometers facilitate compensation in the provided software, but may not provide an option to select a population. 


Functions are provided in the R/bioconductor flowCore package to automate generation of the spillover matrix used to compensate datasets for fluorescent spillover.

For some cytometers, the name of the spillover matrix may be listed as `$SPILLOVER` or `$SPILL`. To determine which keyword to use, the following line will return all of the keywords associated with a sample: `keyword(fs[[1]])`. From there, we can look for keywords like `$SPILL` or `$SPILLOVER.` 

In the case of our dataset cytometer software generates a compensation matrix for you using a wizard or similar algorithm, you can likely extract this from the flowSet and use it to compensate with the following:

In our example dataset, RFP is used to indicate transfected cells, while GFP acts as a reporter gene for our different constructs.


```{r}

### ------------------  Apply Compensation ------------------ ### 

# Retrieve the spillover matrix
keyword(fs[[7]])$SPILL # '$SPILL' is cytometer specific and may need to be updated.

# Apply the spillover matrix to the dataset
fs.comp <- fsApply(fs,function(frame){
     comp <- keyword(frame)$SPILL  # '$SPILL' is cytometer specific and may need to be updated.
     new_frame <- compensate(frame,comp)
     new_frame
})
```

Next we will log transform the data. Log transformation can be used to address skewness in datasets to achieve a near-normal distribution or linearity. Log transformation can make the data easier to interpret as it may make certain relationships more apparent. Some cytometers may have the FCS files already log transformed. Look for a `$TRANSFORM` keyword with `keyword(flowSet[[1]])` to make this determination. Keep in mind that your cytometer may use a slightly different keyword. 

```{r message = FALSE, warning = FALSE}

### ---------------- Log Transform the Compensated Flowset ---------------- ### 

# Log transformation will likely introduce NaNs (not a number) into the data when taking the log of 0. This will return a warning, but is it not an error and they will be excluded in the next step.

# Log transformation of compensated flowset FSC area and height, SSC, red (FL3), and green (FL1) detectors
fs.comp.log <- transform(fs.comp, 
                         `FSC.A` = log(`FSC.A`),
                         `SSC.A` = log(`SSC.A`),
                         `FSC.H` = log(`FSC.H`),
                         `FL1.A` = log(`FL1.A`),
                         `FL3.A` = log(`FL3.A`))


# If you're using the Attune NxT platform use the following lines instead
# fs.comp.log <- transform(fs.comp, 
#                          `FSC.A` = log(`FSC.A`),
#                          `SSC.A` = log(`SSC.A`),
#                          `FSC.H` = log(`FSC.H`),
#                          `BL1.A` = log(`BL1.A`),
#                          `YL1.A` = log(`YL1.A`))

# Exclude NaNs from the compenstated flowset                        
fs.comp.log <- fsApply(x = fs.comp.log, FUN = function(frame){
  values <- exprs(frame)
  exprs(frame) <- na.omit(values)
  frame})

### ------  Log Transform the Uncompensated Flowset (for comparison) ------ ### 

# Log transformation of uncompensated flowset FSC area and height, SSC, red (FL3), and green (FL1) detectors
fs.log <- transform(fs, 
                         `FSC.A` = log(`FSC.A`),
                         `SSC.A` = log(`SSC.A`),
                         `FSC.H` = log(`FSC.H`),
                         `FL1.A` = log(`FL1.A`),
                         `FL3.A` = log(`FL3.A`))


# Exclude NaNs from the uncompensated flowset      
fs.log <- fsApply(x = fs.log, FUN = function(frame){
  values <- exprs(frame)
  exprs(frame) <- na.omit(values)
  frame})
```

## Expression Quantification

To begin quantifying changes in reporter gene expression between our constructs, we first need to identify the live transformed cells based on their expression of red fluorescent protein (RFP) from the pBeaconRFP vector. Dead cells should not fluoresce. In order to identify the transformed fluorescent population of cells, we need a non-transformed control sample to set a threshold of fluorescence. 

We can plot one of our samples on a FL1.A (GFP) vs. FL3.A (RFP) plot to visualize our cell populations. In our example dataset, we can see two distinct cell populations: a GFP positive population, and a population that is positive in both GFP and RFP. There is also a third cluster of events that is likely predominantly debris and non-transformed cells as these events have relatively low GFP and RFP levels.

Note: If your cytometer uses BL1 and YL1 instead of FL1 and FL3, you will have to manually change it throughout the script. You can use Ctrl+F (Windows) or Command+F (Mac) to access the 'find and replace' feature if you are using RStudio. 

In the example dataset, we have overexpressed ARF5Delta, ARF7Delta, and a GUS control using the pBeaconRFP vector and subsequently measured expression from a DR5::GFP reporter. 


```{r message = FALSE, warning = FALSE}

### Plot All Events (Compensated vs Uncompenstated) - Density (RFP vs. GFP) ###

# Initializes graphics device for exporting the plot as a JPEG
# jpeg("GFP_RFP_All_Events_Comp.jpg", width=2000, height=1800, res=300)

# Creating the plot
ggcyto(data = fs.comp.log[8], mapping = aes(x =!!sym(GreenDetector), y = !!sym(RedDetector))) + 
  # Create a density gradient
  stat_density_2d(aes(fill = after_stat(level)), 
                geom = "polygon", bins = 12, n = 200)+
  scale_fill_viridis_c(option = "plasma") +
  # Uses the id column for plot header
  facet_wrap(~id) +  
  # Sets X and Y axis limits
  coord_cartesian(xlim = log(c(1, 0.5e5)), ylim = log(c(1, 2e6))) +
  # Add X and Y axis labels
  labs(x = "GFP Intensity", y = "RFP Intensity") +
  # Define plot graphical style 
  theme_classic() +
  theme(
  panel.border = element_rect(color = "black", fill = NA, size = 1))+
  theme(
    strip.text = element_text(size = 14),      # plot header
    axis.title = element_text(size = 16),      # X and Y axis labels
    axis.text = element_text(size = 12)        # axis tick labels
  )
# dev.off()

# jpeg("GFP_RFP_All_Events_UnComp.jpg", width=2000, height=1800, res=300)
ggcyto(data = fs.log[8], mapping = aes(x =!!sym(GreenDetector), y = !!sym(RedDetector))) + 
  # Create a density gradient
  stat_density_2d(aes(fill = after_stat(level)), 
                geom = "polygon", bins = 12, n = 200)+
  scale_fill_viridis_c(option = "plasma") +
  # Uses the id column for plot header
  facet_wrap(~id) +  
  # Sets X and Y axis limits
  coord_cartesian(xlim = log(c(1, 0.5e5)), ylim = log(c(1, 2e6))) +
  # Add X and Y axis labels
  labs(x = "GFP Intensity", y = "RFP Intensity") +
  # Define plot graphical style 
  theme_classic() +
  theme(
  panel.border = element_rect(color = "black", fill = NA, size = 1))+
  theme(
    strip.text = element_text(size = 14),      # plot header
    axis.title = element_text(size = 16),      # X and Y axis labels
    axis.text = element_text(size = 12)        # axis tick labels
  )
# dev.off()

### -------------- Plot All Events - Dot Plot (RFP vs. GFP) -------------- ### 

## Below will generate the same plots in dot plot format 

# Plot all events of a representative log transformed sample (FL1.A vs. FL3.A)
# jpeg("GFP_RFP_All_Events_Comp_dot.jpg", width=2000, height=1800, res=300)
ggcyto(fs.comp.log[c(8)], mapping = aes( x =!!sym(GreenDetector), y = !!sym(RedDetector))) +
  # Uses the id column for plot header
  facet_wrap(~id) +  
  geom_point(alpha = 0.05, size = 0.5) +
  coord_cartesian(xlim = log(c(1, 0.5e5)), ylim = log(c(1, 2e6))) +
  theme_classic()+
  # Add X and Y axis labels
  labs(x = "GFP Intensity", y = "RFP Intensity") +
  # Define plot graphical style 
  theme(
  panel.border = element_rect(color = "black", fill = NA, size = 1))+
  theme(
    strip.text = element_text(size = 14),      # plot header
    axis.title = element_text(size = 16),      # X and Y axis labels
    axis.text = element_text(size = 12)        # axis tick labels
  )
# dev.off()

# Plot all events of a representative log transformed sample (FL1.A vs. FL3.A)
# jpeg("GFP_RFP_All_Events_UnComp_dot.jpg", width=2000, height=1800, res=300)
ggcyto(fs.log[c(8)], mapping = aes( x = !!sym(GreenDetector), y = !!sym(RedDetector))) +
  # Uses the id column for plot header
  facet_wrap(~id) +  
  geom_point(alpha = 0.05, size = 0.5) +
  coord_cartesian(xlim = log(c(1, 0.5e5)), ylim = log(c(1, 2e6))) +
  # Add X and Y axis labels
  labs(x = "GFP Intensity", y = "RFP Intensity") +
  # Define plot graphical style 
  theme_classic() +
  theme(
  panel.border = element_rect(color = "black", fill = NA, size = 1))+
  theme(
    strip.text = element_text(size = 14),      # plot header
    axis.title = element_text(size = 16),      # X and Y axis labels
    axis.text = element_text(size = 12)        # axis tick labels
  )
# dev.off()
```


## Define a live-cell gate based on positive RFP expression

We can use a control sample with no transformed cells to draw a gate to define the threshold above which truly transformed and thus live cells fall. This is accomplished by drawing a horizontal gate to include only the top 1% of the brightest red fluorescent events on an FL1.A vs. FL3.A plot and exclude 99% of non-fluorescent events. Events that fall above this line in transformed samples will considered to be RFP positive. These RFP-positive events represent viable, transformed cells expressing RFP and the gene of interest from our pBeaconRFP vector. You can adjust the position of this gate by changing value of the `probs` argument in the `gate_quantile()` function. 

```{r message = FALSE, warning = FALSE}
### ------------------  Identify High RFP Events ------------------ ### 

## Identify control sample 
# Get a list of sample names and the corresponding sample # to identify the control sample 
sampleNames(fs.comp.log)

# Assign the # of the control sample from the previous stuff to the 'non_trans_control' variable
non_trans_control <- 4

# Select the top 0.1% of brightest RFP events and assign them to the variable 'RFP_high'
RFP_high <- gate_quantile(fs.comp.log[[non_trans_control]], channel = RedDetector, probs = .995, filterId = "RFP_high") 

### ----------------  Plot RFP Fluorescence Intensity Gate ---------------- ### 

# reorder samples so the control will plot first
# Use the actual values in the column, not just existing factor levels
all_ids <- unique(as.character(pData(fs.comp.log)$id))
other_ids <- setdiff(all_ids, "Non-transformed control_1")

pData(fs.comp.log)$id <- factor(pData(fs.comp.log)$id,
                                levels = c("Non-transformed control_1", sort(other_ids)))

# Initializes graphics device for exporting the plot as a JPEG
# jpeg("RFP_High_All_Events_density.jpg", width=2000, height=1200, res=300)

# Creates a plot of the control sample and a transformed sample (FL1.A vs. FL3.A) with "RFP High" gate
ggcyto(data = fs.comp.log[c(4,8)], mapping = aes(x =!!sym(GreenDetector), y = !!sym(RedDetector))) + 
  # Create a density gradient
  stat_density_2d(aes(fill = after_stat(level)), 
                geom = "polygon", bins = 12, n = 200)+
  scale_fill_viridis_c(option = "plasma") +
  # Uses the id column for plot header
  facet_wrap(~id) +  
  # Sets X and Y axis limits
  coord_cartesian(xlim = log(c(1, 0.5e5)), ylim = log(c(1, 2e6))) +
  # Add X and Y axis labels
  labs(x = "GFP Intensity", y = "RFP Intensity") +
  # Define plot graphical style 
  theme_classic() +
  theme(
  panel.border = element_rect(color = "black", fill = NA, size = 1))+
  theme(
    strip.text = element_text(size = 14),      # plot header
    axis.title = element_text(size = 16),      # X and Y axis labels
    axis.text = element_text(size = 12)        # axis tick labels
  )+
  geom_gate(RFP_high)

# dev.off()

## Creating the same plot in a dot plot format

# jpeg("RFP_High_All_Events_dot.jpg", width=2000, height=1200, res=300)

# Plot the control sample (FL1.A vs. FL3.A) with a horizontal red line delineating the threshold of what is considered an RFP high event.  
ggcyto(fs.comp.log[c(4,8)], mapping = aes( x = !!sym(GreenDetector), y = !!sym(RedDetector))) +  
  facet_wrap(~id) +  
  geom_point(alpha = 0.05, size = 0.5) + 
  coord_cartesian(xlim = log(c(1, 0.5e5)), ylim = log(c(1, 2e6)))+
  labs(x = "GFP Intensity", y = "RFP Intensity") +
  # Define plot graphical style 
  theme_classic() +
  theme(
  panel.border = element_rect(color = "black", fill = NA, size = 1))+
  theme(
    strip.text = element_text(size = 14),      # plot header
    axis.title = element_text(size = 16),      # X and Y axis labels
    axis.text = element_text(size = 12)        # axis tick labels
  )+
  geom_gate(RFP_high)

# dev.off()
```

Next, we will backgate this population of RFP positive viable cells to a FSC.A vs. SSC.A plot to visualize all live cells. To do this, we will plot these 'RFP_high' events on a FSC.A vs SSC.A plot (AKA backgating) to determine where viable cells fall. We can draw a new gate that encompasses 99% of the RFP-positive events and define it as `viable_cells`. This helps us exclude the majority of debris from our analysis.

To define this gate we use `gate_flowclust_2D()`, which uses a robust clustering algorithm to define populations of cells and defines an ellipse centered about that population and encompassing a certain quantile of the events in the population. Decreasing the `quantile` argument will decrease the area of the gate.

```{r message = FALSE, warning = FALSE}

### ---------  Use RFP High Events to Define a Viable Cells Gate  --------- ### 

# Use the RFP high events to define where viable cells fall on a FSC.A vs SSC.A plot
viable_cells <- gate_flowclust_2d(Subset(fs.comp.log[[8]], RFP_high),  
                                  xChannel = "FSC.A", yChannel = "SSC.A",
                                  K = 1, quantile = 0.9)

# Subsetting RFP high events  
fs.sub <- Subset(fs.comp.log, RFP_high)

# Plot RFP_high cells (FSC.A vs SSC.A) and export the plot as a jpg
# jpeg("RFP_high_cells.jpg", width=2000, height=1800, res=300)

ggcyto(fs.sub[c(8)], aes(x = !!FSC_A, y = !!SSC_A)) +
  geom_point(alpha = 0.05, size = 0.5) + 
  facet_wrap(~id) +
  labs(x = "Forward Scatter Area (FSC-A)", y = "Side Scatter Area (SSC-A)" ) +
  theme_classic() +
  theme(
    panel.border = element_rect(color = "black", fill = NA, size = 1),
    strip.text = element_text(size = 14),
    axis.title = element_text(size = 16),
    axis.text = element_text(size = 12)
  ) +
  geom_gate(viable_cells, colour = '#fda53a', size = 2) # Drawing viable cells gate 

# dev.off()

# Plot viable cells against all events on a FSC.A vs SSC.A plot. RFP high events are colored red. 
# jpeg("RFP_high_cells_all.jpg", width=2000, height=1800, res=300)

ggcyto(fs.comp.log[c(8)], mapping = aes( x = !!FSC_A, y = !!SSC_A)) + 
  geom_point(alpha = 0.05, size = 0.5) + 
  facet_wrap(~id) +
  labs(x = "Forward Scatter Area (FSC-A)", y = "Side Scatter Area (SSC-A)" ) +
  theme_classic() +
  theme(
    panel.border = element_rect(color = "black", fill = NA, size = 1),
    strip.text = element_text(size = 14),
    axis.title = element_text(size = 16),
    axis.text = element_text(size = 12)
  ) +
  # Plotting RFP high events on top 
  geom_point(data = fs.sub[[8]], color = 'red', alpha = 0.05, size =   0.5)+
  # Drawing viable cells gate
  geom_gate(viable_cells, colour = '#fda53a', size = 2) 

# dev.off()

```

Next we will screen out any doublets, cells that are stuck together. Cell doublets may artificially inflate measurements, so we want to remove as many doublets as possible without cutting off events/cells from the top of the distribution because those should not be doublets. If you have enough cells, it is generally better to set tighter constraints to minimize inflation of measurements.  We can begin screening out doublet events by first creating a FSC.A vs. FSC.H plot. Proper singlets should have a 1-to-1 correlation between area and height. A doublet will have roughly a 2-to-1 relationship between area and height. By drawing a gate around the events that follow this expected distribution we can exclude the majority of doublet events.  

The default prediction level in the `gate_singlet()` function is set to `0.99` meaning that 1% of cells are estimated to be doublets. You can change the doublet estimation by changing the value assigned to the `prediction_level` argument in the `gate_singlet()` function. In this case we set `prediction_level = 0.9`, which generates a gate excluding the lower tail of cells with FSC-A values greater than FSC-H, and only excluding a small fraction of events in the top right with FSC-H greater than FSC-A. 

```{r message = FALSE, warning = FALSE}
### ------------------  Exclude Cell Doublets  ------------------ ### 

# Subset viable cells   
fs.sub.cells <- Subset(fs.comp.log, viable_cells)

# Plot viable cells (FSC.A vs. FSC.H) and export plot as a .jpg file
# jpeg("Viable_Cell_Population.jpg", width=2000, height=1800, res=300)

ggcyto(fs.sub.cells[8], mapping = aes( x = "FSC.A", y = "FSC.H")) + 
  geom_point(alpha = 0.05, size = 0.5) +
  facet_wrap(~id) +
  labs(x = "Forward Scatter Area (FSC-A)", y = "Side Scatter Area (SSC-A)" ) +
  theme_classic()+
  theme(
    panel.border = element_rect(color = "black", fill = NA, size = 1),
    strip.text = element_text(size = 14),
    axis.title = element_text(size = 16),
    axis.text = element_text(size = 12)
    )+
    coord_cartesian(xlim = c(10.5,14), ylim = c(9.5,13.5))

# dev.off() 

# Subset singlets using FSC.A vs. FSC.H, estimating 5% of cells are doublets. 
singlets <- gate_singlet(Subset(fs.comp.log[[non_trans_control]], viable_cells), 
                         area = "FSC.A", height = "FSC.H", prediction_level = 0.9 )

# Visualize the gate around viable cell singlets (FSC.A vs. FSC.H)
# jpeg("SingletsGate.jpg", width=2000, height=1800, res=300)

ggcyto(fs.sub.cells[8], mapping = aes( x = "FSC.A", y = "FSC.H")) + 
  facet_wrap(~id) +
  geom_point(alpha = 0.05, size = 0.5) + 
  labs(x = "Forward Scatter Area (FSC-A)", y = "Side Scatter Area (SSC-A)" ) +
  theme_classic() +
  theme(
    panel.border = element_rect(color = "black", fill = NA, size = 1),
    strip.text = element_text(size = 14),
    axis.title = element_text(size = 16),
    axis.text = element_text(size = 12)
    ) +
    coord_cartesian(xlim = c(10.5,14), ylim = c(9.5,13.5))+
    geom_gate(singlets, colour = '#0081C7', size = 1) 

# dev.off()

```


Now we have identified the population of viable cells that we want to measure the reporter gene expression in response to our gene of interest.

## Gate Red positive

Now we can finally get to answering the question we were after, “How does overexpression of these transcription factors in the pBeaconRFP vector affect our GFP reporter gene?”

First we will use our gates to select the viable, transformed cells. 

We essentially readjust our gate now that we have excluded the majority of debris with high autofluorescence and thus we can lower our RFP positive threshold. 

```{r message = FALSE, warning = FALSE}
### ------------------  Identify Truly Transformed Cells  ------------------ ### 

# Using the viable cells and singlets gates to subset the cells of interest
fs.single.cells <- Subset(fs.sub.cells, singlets)

# Using the control sample to create a gate of RFP_high cells using only viable singlets
RFP_high <- gate_quantile(fs.single.cells[[non_trans_control]], 
                          channel = RedDetector, probs = .995, 
                          filterId = "RFP_high")

# Initializes graphics device for exporting the plot as a JPEG
# jpeg("TransformedCells_Gate_Density.jpg", width=2000, height=1000, res=300)

# Creating the plot
ggcyto(data = fs.single.cells[c(4,8)], mapping = aes(x =!!sym(GreenDetector), y = !!sym(RedDetector))) + 
  # Create a density gradient
  stat_density_2d(aes(fill = after_stat(level)), 
                geom = "polygon", bins = 12, n = 200)+
  scale_fill_viridis_c(option = "plasma") +
  # Uses the id column for plot header
  facet_wrap(~id) +  
  # Sets X and Y axis limits
  coord_cartesian(xlim = log(c(1, 0.5e5)), ylim = log(c(1, 2e6))) +
  # Add X and Y axis labels
  labs(x = "GFP Intensity", y = "RFP Intensity") +
  # Define plot graphical style 
  theme_classic() +
  theme(
  panel.border = element_rect(color = "black", fill = NA, size = 1))+
  theme(
    strip.text = element_text(size = 14),      # plot header
    axis.title = element_text(size = 16),      # X and Y axis labels
    axis.text = element_text(size = 12)        # axis tick labels
  ) + 
  geom_gate(RFP_high, colour = '#FF4251', size = 2)

# dev.off()


## Below will generate the same plot in dot plot format 

# Plot all events of a representative log transformed sample (FL1.A vs. FL3.A)
# jpeg("TransformedCells_dot.jpg", width=2000, height=1250, res=300)

ggcyto(fs.single.cells[c(4,8)], mapping = aes( x =!!sym(GreenDetector), y = !!sym(RedDetector))) +
  # Uses the id column for plot header
  facet_wrap(~id) +  
  geom_point(alpha = 0.05, size = 0.5) +
  coord_cartesian(xlim = log(c(1, 0.5e5)), ylim = log(c(1, 2e6))) +
  theme_classic()+
  # Add X and Y axis labels
  labs(x = "GFP Intensity", y = "RFP Intensity") +
  # Define plot graphical style 
  theme(
  panel.border = element_rect(color = "black", fill = NA, size = 1))+
  theme(
    strip.text = element_text(size = 14),      # plot header
    axis.title = element_text(size = 16),      # X and Y axis labels
    axis.text = element_text(size = 12)        # axis tick labels
  )+
  geom_gate(RFP_high, colour = '#FF4251', size = 2)

# dev.off()

### ------------------  Subset Truly Transformed Cells ------------------ ### 

# Subset transformed viable signlets
fs.transformed.cells <- Subset(fs.single.cells, RFP_high)
```

We can also now calculate the transformation efficiency, which is the number of transformed cells divided by the total number of cells. 

```{r message = FALSE, warning = FALSE}
### ----------------- Calculate Transformation Efficiency ----------------- ### 

# Calculate transformation efficiency for each construct 
percent_transformed <- lapply(sampleNames(fs.single.cells), function(s) {
  total_events <- nrow(exprs(fs.single.cells[[s]]))
  transformed_events <- nrow(exprs(Subset(fs.single.cells[[s]], RFP_high)))
  construct_label <- pData(fs.single.cells)[s, "construct"]
  data.frame(
    sample = s,
    construct = construct_label,
    total = total_events,
    transformed = transformed_events,
    percent_transformed = round(100 * transformed_events / total_events, 2)
  )
}) %>% bind_rows()

# Calculate the mean and SE transformation rates 
summary_df <- percent_transformed %>%
  group_by(construct) %>%
  summarise(
    mean_percent = mean(percent_transformed),
    se = sd(percent_transformed) / sqrt(n())
  )
# Define sample order for plotting
summary_df$construct <- factor(summary_df$construct, levels = c(
  "Non-transformed control",
  "GUS",
  "GUS + IAA",
  "ARF7"
))

# Plot transformation efficiency and export as a JPG
# jpeg("TransformationRates.jpg", width=1250, height=2250, res=300)

ggplot(summary_df, aes(x = construct, y = mean_percent, fill = construct)) +
  geom_col(width = 0.6) +  # bar chart
  geom_errorbar(aes(ymin = mean_percent - se, ymax = mean_percent + se),
                width = 0.2, size = 1, color = "gray10") +  # error bars
  scale_fill_manual(values = c(
    "GUS + IAA" = "#fda732",
    "ARF7" = "#ef864d",
    "GUS" = "#f9d02f",
    "Non-transformed control" = "#555555"
  )) +
  labs(
    x = "",
    y = "Transformation Efficiency (%)",
  ) +
  theme_classic()+
  theme(
    legend.position = "none",
    panel.border = element_rect(color = "black", fill = NA, size = 1),
    axis.text.x = element_text(angle = 45, hjust = 1, size = 15),
    axis.title.y = element_text(size = 15),
    )

# dev.off()

### ------------------  Plot Only Truly Transformed Cells ------------------ ### 

## Plot the final subsetted flowset 
# Initializes graphics device for exporting the plot as a JPEG
# jpeg("Subset_TransformedCells.jpg", width=2000, height=1800, res=300)

# Creating the plot
ggcyto(data = fs.transformed.cells[c(8)], mapping = aes(x =!!sym(GreenDetector), y = !!sym(RedDetector))) + 
  # Create a density gradient
  stat_density_2d(aes(fill = after_stat(level)), 
                geom = "polygon", bins = 12, n = 200)+
  scale_fill_viridis_c(option = "plasma") +
  # Uses the id column for plot header
  facet_wrap(~id) +  
  # Sets X and Y axis limits
  coord_cartesian(xlim = log(c(1, 0.5e5)), ylim = log(c(1, 2e6))) +
  # Add X and Y axis labels
  labs(x = "GFP Intensity", y = "RFP Intensity") +
  # Define plot graphical style 
  theme_classic() +
  theme(
  panel.border = element_rect(color = "black", fill = NA, size = 1))+
  theme(
    strip.text = element_text(size = 14),      # plot header
    axis.title = element_text(size = 16),      # X and Y axis labels
    axis.text = element_text(size = 12)        # axis tick labels
  ) 
# dev.off()

```

```{r}
### ------------------  Convert Flowset to a Dataframe ------------------ ### 

# Convert flowset into a dataframe
data <- steadyState(fs.transformed.cells, gated = TRUE)

# Remove the non-transformed sample from the dataset
data <- subset(data, !grepl("Non-transformed control", construct))
  
#Reordering the samples for plotting
relevels <- c("GUS", "GUS + IAA", "ARF7") 
data$construct <- factor(data$construct, levels = relevels)

```


```{r message = FALSE, warning = FALSE}

# Create a variable that uniquely identifies each replicate
data <- data %>%
  mutate(
    replicate_id = paste0(construct, "_", replicate),
    construct = fct_rev(as.factor(construct))
    )

# Defining replicate and sample colors to use for plotting. The bottom color for each sample will be plotted on top. 
replicate_colors <- c(
  "GUS_1" = "#EA7750",
  "GUS_2" = "#FF9933",
  "GUS_3" = "#ffde55",
  "GUS + IAA_1" = "#EA7750",
  "GUS + IAA_2" = "#ffde55",
  "GUS + IAA_3" = "#FF9933",
  "ARF7_1" = "#ffde55",
  "ARF7_2" = "#FF9933",
  "ARF7_3" = "#EA7750"
)

# jpeg("Density_Ridges.jpg", width = 1750, height = 1000, res = 300)

ggplot(data, aes(y = construct)) +
  geom_density_ridges(
    aes(x = as.numeric(!!sym(GreenDetector)),
      fill = replicate_id),
    color = "black",     # solid black outlines
    from = 0,
    to = 11.5
  ) +
  scale_fill_manual(values = alpha(replicate_colors, .7)) +
  scale_y_discrete(expand = expansion(add = c(0, 2))) +
  labs(x = "Log10 GFP Intensity", y = "") +
  theme_classic() +
  theme(
    axis.text.y = element_text(size = 9),
    legend.position = "none",
    panel.border = element_rect(color = "black", fill = NA, size = 1))

# dev.off()
```

## Calculate Summary Statistics

Then we can calculate summary statistics for each sample.

We can now run an ANOVA on our samples to determine the probability that any differences we observe are due to chance. 

```{r summary stats}


replicate_means <- data %>%
  group_by(construct, replicate) %>%
  summarise(mean_signal = mean(!!sym(GreenDetector)), .groups = "drop")

data.aov <- aov(mean_signal ~ construct, data = replicate_means)
summary.aov <- summary(data.aov)
summary.aov


leveneTest(mean_signal ~ construct, data = replicate_means)
# Q-Q plot of residuals

# jpeg("QQplot.jpg", width = 2000, height = 2000, res = 300)
qqnorm(residuals(data.aov))
qqline(residuals(data.aov), col = "blue")

# dev.off

durbinWatsonTest(data.aov)


data.hsd <- HSD.test(data.aov, trt = "construct", alpha = 0.01)
data.hsd


```

Next we can conduct a Tukey HSD post-hoc analysis to group samples that are statistically similar to each other.

```{r}
# Create a new data frame with the average FL1.A activity for each construct and its associated HSD group 
HSD.groups <- 
data.hsd$groups %>% as_tibble(rownames = "names") %>%
  mutate(construct = names) %>% 
  mutate(FL1.A = mean_signal) %>% 
  left_join(data.hsd$means %>% 
              as_tibble(rownames = "names") %>% 
              dplyr::select(c(names)), 
            by = "names") %>%
  mutate(construct = fct_relevel(construct, "GUS", after = 0))
```

## Generate Figures

With all the processing and statistics completed, we can finally get to plotting our results. We first plot our data using a violin boxplot. This offers a comprehensive view of the data distributions along with the summary statistics. 

```{r}
# Reordering the samples for plotting
data$construct <- factor(data$construct, levels = relevels)

# Plot data with a violin boxplot. mean = green, median = black
# jpeg("Violin_Boxplot2.jpg", width = 3000, height = 3000, res = 500)

ggplot(data,
  aes(y = as.numeric(FL1.A), x = construct, fill = construct)) + 
  geom_violin(width = .75, color = "gray20", size=.7) +
  geom_boxplot(width = 0.25, color = "gray20", fill = 'white', size = .7, fatten = 2) +
  stat_summary(fun = mean, geom = "crossbar", width = 0.24, color = "#AFC247", size = 0.5) +
  scale_fill_manual(values = c("#f9d02f", "#fda732", "#ef864d")) +  # custom colors here
  theme(legend.position = "none") +
  ylab("Log10 GFP Intensity") +
  xlab(" ") +
  geom_text(data = HSD.groups, aes(label = groups), size = 6, vjust = -6, hjust = 0.5) +
  theme_classic()+
  theme(
    legend.position = "none",
    axis.text.x = element_text(angle = 45, hjust = 1, size = 15),
    axis.title.y = element_text(size = 15),
    panel.border = element_rect(color = "black", fill = NA, size = 1))+
  coord_cartesian(ylim = c(-6, 13))
# dev.off()

```


```{r}

# Calculating fold change by dividing the value of the experimental group by the value of the control group
#undoing log transformation
data.exp <- transform(data, 
                         `FSC.A` = exp(`FSC.A`),
                         `SSC.A` = exp(`SSC.A`),
                         `FSC.H` = exp(`FSC.H`),
                         `FL1.A` = exp(`FL1.A`),
                         `FL3.A` = exp(`FL3.A`))

# Calculate the mean FL1.A for the control group
control.mean <- data.exp %>%
  dplyr::filter(construct == "GUS") %>%
  summarize(gus_value = mean(FL1.A)) %>%
  pull(gus_value)

# Take the mean DR5:GFP activity of all samples
data.mean <- aggregate(FL1.A ~ construct + replicate, data = data.exp, FUN = mean)

# Calculating average fold of DR5:GFP from control group
data.mean$FoldChange <- (data.mean$FL1.A / control.mean) 

# Calculate the standard error for each construct
data.se <- data.mean %>%
  group_by(construct) %>%
  summarize(SE = sd(FoldChange) / sqrt(n()))


data.avg <- data.mean %>%
  group_by(construct) %>%
  summarize(FoldChange = mean(FoldChange))

# Merging with standard error
data.avg <- data.avg %>%
  left_join(data.se, by = "construct") 


letters_df <- data.hsd$groups %>%
  as_tibble(rownames="construct") %>%
  dplyr::select(construct, groups)

data.final <- data.avg %>%
  left_join(letters_df, by="construct")

# Reordering the samples for plotting
data.final$construct <- factor(data.final$construct, levels = relevels)

# Plotting the data
# jpeg("Relative_Change.jpg", width = 1250, height = 2500, res = 500) 

ggplot(data.final, aes(x = construct, y = FoldChange), fill = construct) +
  geom_bar(stat = "identity", 
           width = 0.5, 
           # color = "black", # for outlines
           fill = c("#f9d02f", "#fda732", "#ef864d")) + 
   geom_errorbar(aes(ymin = FoldChange - SE,
                    ymax = FoldChange + SE),
                width = 0.25,  # Adjust the width of the error bars
                size = .75,
                color = "grey7") +  # Error bar color
  geom_text(aes(label = FoldChange %>% round(2)),
            size = 4, 
            position = position_nudge(y = 0.15),
            vjust = ifelse(data.final$FoldChange >= 0, -0.5, -3.5)) +
  geom_text(data = data.final,
            aes(label = groups,
            x = construct),
            size = 5.5,
            position = position_nudge(y = .3),
            vjust = ifelse(data.final$FoldChange >= 0, -1, -3.5),
            color = "grey7") +
  geom_hline(yintercept = 0, 
              color = "gray15", 
              size = 0.5) + 
  ylab("Fold change \n GFP activation from control") + 
  xlab(" ") + 
theme(
    panel.border = element_rect(color = "black", fill = NA, size = 1),
    panel.background = element_blank(),  # Remove panel background
    plot.background = element_blank(),  # Remove plot background
    panel.grid.major = element_blank(), # Remove major gridlines
    panel.grid.minor = element_blank(), # Remove minor gridlines
    axis.text.x = element_text(angle = 45, hjust = 1, size = 10),  # Rotate x-axis labels
  ) +
  coord_cartesian(ylim = c(0, 2.7))+
  scale_y_continuous(breaks = seq(0, 10, by = 0.5))

# dev.off()


```
````
